# Gene Therapy with the N-terminal Fragment of Na_v_1.5 for Cardiac Channelopathies: A Novel Transcomplementation Mechanism Potentiating the Cardiac Sodium Current

**DOI:** 10.1101/2024.10.30.621028

**Authors:** Marie Gizon, Marine C. Ferrand, Vincent Fontaine, Nathalie Mougenot, Pierre-Léo Laporte, Nathalie Gaborit, Fabrice Extramiana, Isabelle Baró, Pascale Guicheney, Flavien Charpentier, Nicolas Doisne, Nathalie Neyroud

## Abstract

**BACKGROUND:** Cardiac channelopathies, caused by mutations in ion-channel genes, can lead to sudden cardiac death (SCD) *via* ventricular arrhythmias. Brugada syndrome (BrS) is a rare inherited channelopathy characterized by a unique ECG pattern and a high incidence of ventricular fibrillation leading to SCD in the absence of structural heart defects. The main gene responsible for 20-25% of BrS cases is *SCN5A*, encoding the cardiac sodium channel α-subunit Na_v_1.5, which carries the sodium current (*I*_Na_) responsible for the rapid depolarization phase of the action potential (AP). While current treatments do not target the genetic cause of channelopathies, this study explores the therapeutic potential of overexpressing the N-terminal region of Na_v_1.5 (Nter) to restore electrical activity by rescuing *I*_Na_, in the context of *SCN5A* deficiency.

**METHODS:** We overexpressed the Nter peptide using viral vectors in *Scn5a*^+/-^ mice, in CRISPR/Cas9 edited-*SCN5A*^+/-^ cardiomyocytes derived from induced-pluripotent stem cells (iPSC-CMs) and in BrS patient iPSC-CMs. We assessed molecular and functional effects of Nter overexpression *in vitro* and *in vivo* by measuring Na_v_1.5 subcellular expression and electrophysiological activity and by recording ECGs and arrhythmias.

**RESULTS:** Whereas *Scn5a*^+/-^ mice showed an impaired *I*_Na_ associated with a slowed-cardiac conduction characteristic of the BrS phenotype, cardiac-specific overexpression of Nter corrected AP parameters by restoring *I*_Na_ density in *Scn5a^+/-^* mouse cardiomyocytes. This increase in *I*_Na_ density was caused by a translocation of Na_v_1.5 to the cell membrane in Nter-overexpressing mice. Most importantly, Nter overexpression normalized atrioventricular and ventricular conduction and protected *Scn5a*^+/-^ mice from arrhythmias triggered by programmed electrical stimulation. Similarly, Nter overexpression in *SCN5A*^+/-^ human iPSC-CMs led to a 2-fold increase in Na_v_1.5 cell-surface expression, resulting in normalization of *I*_Na_ and AP parameters and abolition of early after depolarizations observed during spontaneous AP recordings. Similar results were obtained in iPSC-CMs derived from a BrS patient, confirming the potential of this therapy in human models.

**CONCLUSIONS:** This study identified a novel therapeutic peptide effective in restoring cardiac excitability in animal and cellular models of BrS, paving the way for future development of therapies for life-threatening arrhythmias in patients with *SCN5A* deficiency.

## INTRODUCTION

Cardiac channelopathies result from genetic mutations predisposing to a higher susceptibility to life-threatening arrhythmias associated with an increased lifetime risk of sudden cardiac death (SCD). Although they account for a small proportion of all SCDs, these disorders are responsible for the majority of SCDs in young people with otherwise normal hearts and life expectancy^1^. Of these, Brugada syndrome (BrS) accounts for 20-25% of SCDs occurring in the absence of any apparent structural heart abnormality^2^. To date, a causal genetic variant has been identified in ≍35% of BrS probands in one of the >23 genes that have been related to BrS. However, the majority of these variants have been discovered by candidate gene approach in few cases and are linked to <5% of total BrS cases, questioning their role in BrS pathophysiology^3^. In addition, recent genome-wide association studies have reported common variants in susceptibility genes, that may explain the incomplete penetrance and phenotypic heterogeneity of BrS, suggesting a polygenic origin^4,5^. Within the genetic complexity of BrS, *SCN5A* is nevertheless the only gene considered causative and clinically relevant^6^, as ≍75% of the identified BrS mutations are located in this gene^7^. *SCN5A* encodes the cardiac sodium channel α-subunit Na_v_1.5, which generates the sodium current (*I*_Na_) responsible for the rapid depolarization phase of the action potential (AP)^8^. Of the BrS-pathogenic variants in *SCN5A*, ≍40-50% are truncating mutations, including frameshift, nonsense and splice variants, resulting in a premature stop signal, making haploinsufficiency an important mechanism of BrS (https://www.ncbi.nlm.nih.gov/clinvar and^9^).

According to current clinical guidelines^1,10^, SCD prevention relies on implantable cardioverter defibrillators (ICD), which are associated with a high proportion of long-term complications^11^. Pharmacological treatments are recommended for arrhythmic storms or inappropriate electrical shocks, but do not fully protect against SCD. Epicardial ablation, indicated for patients with recurrent ICD shocks refractory to drug therapy, is invasive and requires long-term risk assessment. In this context, gene therapy directly targeting the arrhythmogenic substrate would be a major breakthrough^12^. To overcome the difficulties and complications associated with adeno-associated virus (AAV)-mediated overexpression of a large gene such as *SCN5A*^13,14^, we hypothesized that using a short fragment of the cardiac sodium channel Na_v_1.5, which we and others have shown to increase *I*_Na_ *in vitro*^15,16^, would restore the electrophysiological properties of cardiomyocytes and thereby normalize the BrS phenotype associated with *SCN5A* haploinsufficiency.

Here, we evaluated the therapeutic potential of the N-terminal region (Nter) of Na_v_1.5 *in vivo* in *Scn5a*^+/-^ mice^17^ and *in vitro* in human induced pluripotent stem cell-derived cardiomyocytes (iPSC-CMs) genome-edited to recapitulate *SCN5A* haploinsufficiency^18^. Our data demonstrated that viral vector-mediated overexpression of Nter increased the cell-surface expression of Na_v_1.5 and restored *I*_Na_ in both models, leading to a normalization of AP and ECG parameters and protected mice from arrhythmias triggered by intracardiac programmed electrical stimulation. Importantly, the Nter gene therapy was also shown to be efficient in iPSC-CMs derived from a BrS patient. Altogether, our results highlight the potential interest of Nter-gene therapy for the treatment of patients with haploinsufficient mutations in *SCN5A* and, more broadly, for patients with cardiac disorders associated with a deficient cardiac sodium channel.

## METHODS

Detailed materials and methods used in this study are included in the Supplemental Material.

### *Scn5a*^+/-^ murine model and AAV injection

All animal experiments complied with the Guide for the Care and Use of Laboratory Animal, according to the directive 2010/63/EU of the European Parliament and were approved by the local committee of animal care (agreement: B751320). Initially created on the 129Sv/Ev genetic background^17^, *Scn5a*^+/-^ mice were transferred on the C57BL/6J background. Three to 4 days after birth, adeno-associated virus serotype 9 carrying the Nter sequence (AAV-Nter) (3.10^12^ viral particles per mouse) and control AAV (AAV-GFP) (2.8 10^11^ viral particles per mouse) were directly injected into the jugular vein of mice as previously described^14^.

### Electrocardiography (ECG) and programmed electrical stimulation

Before performing programmed electrical stimulation (PES), surface ECG was recorded from 9-week-old mice anesthetized with etomidate (*12 mg/kg*) using 29G subcutaneous electrodes connected to the IOX analog-digital converter (*emka Technologies*). ECG parameters (heart rate and P-wave, PR interval and QRS complex durations) were analyzed with ecgAUTO software (*emka Technologies*) on 3-minute basal ECG recordings. Recordings were filtered at 50 Hz before analysis and ECG traces were averaged.

PES was then conducted to determine ventricular refractory period and arrhythmia susceptibility as previously described^19^. The detailed procedure is included in the Supplemental Material.

### Isolation of mice ventricular cardiomyocytes

Cardiomyocytes were isolated using a simplified, Langendorff-free method published in 2016^20^. Briefly, 9-10-week-old mice were anesthetized (*sodium pentobarbital 150 mg/kg*) before chest opening. After cutting the descending aorta and inferior vena cava, EDTA buffer (130 mM NaCl, 5 mM KCl, 0.5 mM NaH_2_PO_4_, 10 mM HEPES, 10 mM Glucose, 10 mM Butanedione-monoxime (BDM), 10 mM Taurine and 5 mM EDTA) was injected into the apex of the right ventricle. The aorta was then clamped, and the heart was removed from the chest and bathed in EDTA buffer. After injection of EDTA buffer into the apex of left ventricle, the perfusion buffer (130 mM NaCl, 5 mM KCl, 0.5 mM NaH_2_PO_4_, 10 mM HEPES, 10 mM Glucose, 10 mM BDM, 10 mM Taurine and 1 mM MgCl2) was injected. Then, collagenase buffer (collagenase 2 (*Worthington*) concentration was depending on the activity) was injected into the apex of the left ventricle until digestion was apparent. The clamp was removed, and the tissue was gently cut into ≍1 mm^3^ pieces and gently flushed with Pasteur pipette to separate cardiomyocytes. Stop buffer (perfusion buffer containing 5% of bovine serum albumin) was then added to stop digestion and the cell suspension was passed through a 100-µm filter. Cardiomyocytes were used for patch-clamp recordings within 5 h.

### Electrophysiological recordings in mice ventricular cardiomyocytes

Sodium currents were recorded by whole-cell voltage clamp technique at room temperature (22 ± 1°C) using Amplifier VE-2 (*Alembic*) as previously described^14^. Currents were filtered at 5 kHz (23 dB, 8-pole low-pass Bessel filter) and digitized at 30 kHz (NI PCI-6251, *National Instruments, Austin, TX, USA*) and processed by the Elphy software (*Gerard Sadoc, CNRS, Gif/Yvette, France*). Sodium current analysis was performed using Sigma plot (*Systat Software, San Jose, USA*). Patch-clamp pipettes (resistance 1 to 1.5 MΩ) (*World Precision Instrument*) were filled with solution composed of (in mM) 135 CsCl, 1 CaCl_2_, 2 MgCl_2_, 4 Mg-ATP, 15 EGTA, 10 HEPES adjusted to pH 7.2 with CsOH and cells were superfused with an external solution containing (in mM) 10 NaCl, 123.5 CsCl, 2 CaCl_2_, 2.5 MgCl_2_, 10 HEPES, 10 glucose, 20 Tetra Ethyl Ammonium (TEA), 3 4-AminoPyridine (4-AP), 3 CoCl_2_ adjusted to pH 7.4 with CsOH.

To determine peak *I*_Na_ amplitude and current-voltage relationship (I/V curves), currents were elicited by test potentials of 0.2 Hz frequency from -100 to +60 mV for 50 ms by increments of 10 mV from a holding potential of -120 mV. The currents (pA) were normalized by the membrane capacitance (pF), which reflects the cell size. Sodium channel activation was fitted using the Boltzmann equation: G = G_max_/(1 + exp(V_1/2_ - V_m_)/k). Steady-state of inactivation was established with the following protocol: from a holding potential of -120 mV, pre-pulses of 2 s were applied from -140 to +20 mV with 10-mV increments followed by a 50-ms test pulse to -20 mV at 0.2 Hz frequency. The Boltzmann equation, I =I_max_/(1 + exp(V_1/2_ - V_m_)/k) was applied to fit the inactivation curve.

APs were recorded using the standard microelectrode technique on the right ventricular free wall of 9 to 10-week-old mice. The preparations were stimulated (Isostim A320, *World Precision Instruments*) *via* a bipolar Ag-electrode with 1-ms pulses at twice the threshold voltage. APs were recorded using 15-30 MΩ standard glass microelectrodes filled with 3 M KCl and coupled to a high-input impedance preamplifier (Intracellular Electrometer IE-210, *Warner Instrument Corp*). APs were displayed on an oscilloscope and simultaneously digitized and analyzed with the interactive Acquis1 software, version 4.0 (*Gerard Sadoc, CNRS, Gif/Yvette, France*). Preparations were stimulated at 1 Hz and allowed to equilibrate for ≥1 h, before the onset of recording. AP measurements included maximum upstroke velocity (dV/dt_max_) and AP duration (APD) at 5, 50 and 90 % (APD5, 50, 90) of repolarization.

### Differentiation and adenoviral transduction of iPSC-CMs

The iPSCs were cultured on Matrigel-coated plates in mTESR™1 media (*STEMCELL Technologies*) and passaged manually every 4 to 6 days. Cardiac differentiation of iPSCs was carried out according to a previously published protocol based on sequential application of small molecules modulating the Wnt signaling pathway^21^. Briefly, iPSCs were dissociated using Gentle cell reagent (*STEMCELL Technologies*) and plated in mTeSR®1 with 10 μM Y-27632 (*Biotechne*) for 24 h and cultured in mTeSR®1 until the cells reached 80-90 % confluence. The differentiation was then initiated by adding 6 μM CHIR99021 (*Abcam*) in RPMI1640 medium (*Gibco*) supplemented with B27 minus insulin (*Gibco*). Two days later, the medium was replaced by RPMI1640 medium supplemented with B27 minus insulin and 5 μM IWP2 (*Tocris Biosciences*). On day 5, cells were cultured in RPMI1640 medium with B27 minus insulin and from day 7, in RPMI1640 medium with complete B27. Presence of beating cells was considered the primary feature of successful cardiac differentiation. All experiments were done using iPSC-CMs from at least 3 independent differentiations.

At differentiation day 20, cardiomyocytes were dissociated using the Multi-dissociation kit 3 (*Miltenyi Biotec*) and sorted using the Pluripotent Stem-Cell (PSC)-derived Cardiomyocyte Isolation kit (*Miltenyi Biotec*) according to manufacturer’s protocols. Cells were then reseeded onto Matrigel-coated plates for adenoviral transduction realized one week later. After 2 washes with Phosphate Buffer Saline (PBS), iPSC-CMs were incubated at 37°C for 2 h with Nter adenovirus (5.10^6^ pi/mL) or with GFP adenovirus (2.10^7^ pi/mL) in serum-free DMEM (*Gibco*).

### Electrophysiological recordings in iPSC-CMs

All patch-clamp recordings were performed in spontaneously beating iPSC-CMs at 3-5 days post-adenoviral transfection, *i.e.* 30-32 days post-differentiation, at 37°C. Data were acquired using Axopatch 200B (*Molecular devices*), processed by the Axon digidata 1550A/pClamp v10.7 acquisition system (*Molecular devices*) and analyzed using Clampfit v10.7 software (*Molecular devices*).

Sodium currents were recorded by whole-cell voltage-clamp. Patch-clamp pipettes (resistance 1-2 MΩ) (*World Precision Instrument*) were filled with a solution containing in mM: 5 NaCl, 130 CsCl, 1 CaCl_2_, 15 HEPES, 10 Glucose, and 3 Mg-ATP adjusted to pH 7.2 with CsOH. Cells were bathed in the extracellular solution consisting of (in mM): 50 NaCl, 90 CsCl, 2 CaCl_2_, 3 MgCl_2_, 10 HEPES and 10 Glucose, adjusted to pH 7.4 with CsOH. Currents were filtered at 5 kHz and digitized at 50 kHz. To determine peak *I*_Na_ amplitude and I/V curves, currents were elicited by test potentials of 0.2 Hz frequency from a holding potential of -120 mV by applying 50-ms depolarizing pulses from -80 to +40 mV by 10-mV increments. The currents (pA) were normalized by the membrane capacitance (pF). From the I/V curves, Na^+^-channel activation was calculated and fitted according to the Boltzmann equation. Na^+^-channel steady-state of inactivation was assessed by applying depolarizing pre-pulses of 500 ms from -130 to -20mV in 10-mV increments followed by a 50-ms test pulse at-20 mV. The Boltzmann equation was applied to fit the inactivation curve.

APs were recorded by the perforated whole-cell current-clamp technique. Patch-clamp pipettes (resistance 1-2 MΩ) (*World Precision Instrument*) were filled with a solution containing in mM: 10 NaCl, 20 KCl, 120 K-gluconate, 2 MgCl_2_, 10 HEPES, and 0.22 Amphotericin adjusted to pH 7.2 with KOH. Cells were bathed in the extracellular solution consisting of (in mM): 140 NaCl, 5.4 KCl, 2 MgCl_2_, 2 CaCl_2_, 10 HEPES and 20 Glucose, adjusted to pH 7.4 with NaOH. The membrane voltage was artificially lowered and maintained at -80 mV by the injection of a holding current during the experiment. APs were induced by a rectangular pulse of 250 to 1500 nA for 1 ms at a frequency of 1.25 Hz.

## Statistical analysis

All statistical analyses were performed using Prism8 v8.3.0 software (*GraphPad*). Data were expressed as means ± SEM or median ± interquartile, as mentioned. After Shapiro-Wilk test to verify the normality of the distribution, Student t test or Mann-Whitney test were used to compare difference between two groups. Kruskal-Wallis test or one-way analysis of variance (ANOVA) with Tukey post-hoc test was applied to compare multiple groups, as appropriate. The Fisher one-tail test was used to compare occurrence of EADs in spontaneous APs.

## RESULTS

### AAV-mediated Nter expression in *Scn5a*-defficient mice was cardiac speciffic

To assess the potential benefit of Nter cardiac overexpression, we used an AAV-systemic-injection strategy^14^ targeting the heart of the *Scn5a*-deficient murine model *Scn5a*^+/-^, which is known to express a BrS-like phenotype^17^. We first determined the safety of AAV delivery in control and *Scn5a*^+/-^ mice cardiac structure and function by performing echocardiography (Table S1). No significant differences were observed in the heart of mice injected with control AAV (AAV-GFP) compared to non-injected mice, confirming that AAV vectors and GFP expression did not alter cardiac function. Similarly, AAV-mediated cardiac overexpression of the Nter (AAV-Nter) did not alter mice echocardiographic features (Table S2).

We then evaluated the cardiac specificity of Nter overexpression by quantifying GFP transcripts in heart, liver, skeletal muscle, and brain tissues (Figure S1A) and AAV transduction efficiency was analyzed by immunochemistry in injected-mice heart tissue (Figure S1B). Our results revealed that combination of the cardiotropism of AAV serotype 9 and the cardiac specificity of the chicken Troponin-T (cTnT) promoter guaranteed a specific and robust cardiac transduction of ≈70% of cells in injected mice. In all further experiments, cardiac transduction efficiency was systematically quantified by RT-qPCR analysis of GFP and Nter expression levels (Figure S1C and S1D). Mice exhibiting GFP expression levels below the lowest level observed in non-injected mice (background signal) were considered not properly injected and were excluded from the study.

### Nter rescued *I*_Na_ in *Scn5a*^+/-^ cardiomyocytes by modifying Na_v_1.5 distribution

Given the role established *in vitro* of the Nter peptide in modulating the cardiac sodium current^15,16^, we sought to decipher its effects *in vivo* in the context of *Scn5a* haploinsufficiency by conducting an electrophysiological study. Whole-cell patch-clamp recordings of *I*_Na_ confirmed a significant decrease in *I*_Na_ peak amplitude in cardiomyocytes isolated from *Scn5a*^+/-^ mice hearts compared to control mice, validating the model of *Scn5a* haploinsufficiency. Most importantly, Nter overexpression in *Scn5a*^+/-^ mice hearts normalized *I*_Na_ density to control levels (Figure 1A) without affecting its biophysical properties (Table 1). We then recorded action potentials (APs) from mice right-ventricle free wall and observed that Nter overexpression significantly restored AP maximum upstroke velocity (dV/dt_max_), which was initially decreased by *Scn5a* haploinsufficiency (Figure 1B). AP duration (APD) shortening observed in *Scn5a*^+/-^ mice hearts was not modified by Nter overexpression (Figure S2).

**Figure 1.**
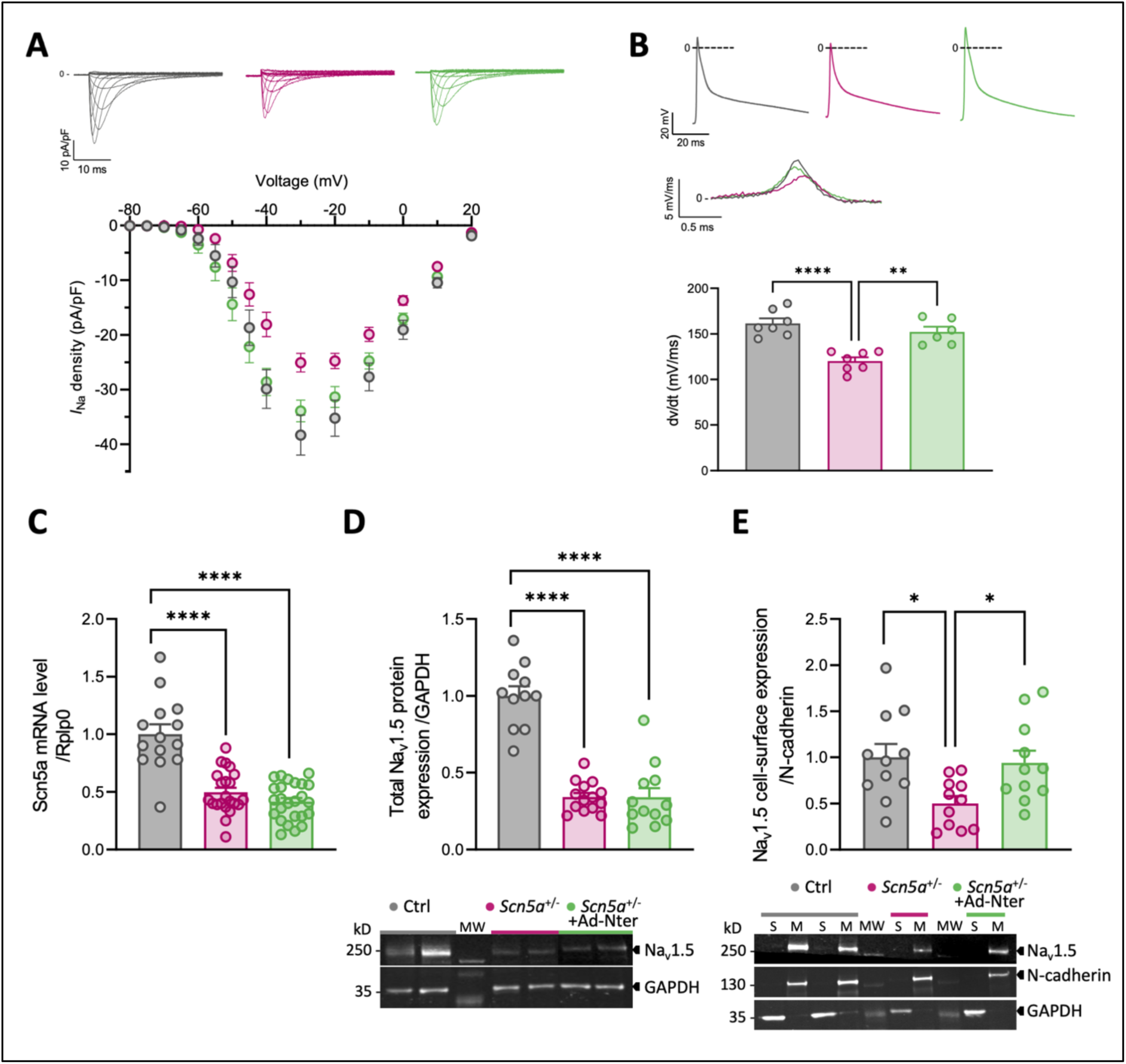
Nter overexpression restored *I*_Na_ density and AP parameters by increasing Na_v_1.5 cell-surface expression in cardiomyocytes of *Scn5a*-defficient mice. **A, Top**. Representative *I*_Na_ recordings in cardiomyocytes from control mice (n= 16) and *Scn5a*^+/-^ mice non-injected (n= 14) or injected with AAV-Nter (n= 15); **Bottom.** Normalized *I*_Na_ density-membrane potential relationships recorded in control (Ctrl) cardiomyocytes (n= 16), *Scn5a*^+/-^ cardiomyocytes (n= 14) and *Scn5a*^+/-^ cardiomyocytes + AAV-Nter (n= 15). **B, Top.** Representative AP and maximum upstroke velocity (dV/dt_max_) traces recorded in Ctrl cardiomyocytes, *Scn5a*^+/-^ cardiomyocytes and *Scn5a*^+/-^ cardiomyocytes + AAV-Nter; **Bottom.** Measurements of dV/dt_max_ in cardiomyocytes of Ctrl and *Scn5a*^+/-^ mice non-injected or injected with AAV-Nter; \*\**P*<0.01, \*\*\*\**P*<0.0001 (*One-way Anova test*). **C.** Scn5a mRNA expression level measured in cardiomyocytes of Ctrl and *Scn5a*^+/-^ mice non-injected or injected with AAV-Nter. Cycle thresholds (Ct) were normalized to *Rplp0* and the ratio *vs.* Ctrl (2^-ΔΔCt^) was then calculated; \*\*\*\**P*<0.0001 (*One-way Anova test*). **D, Bottom.** Western-blot analysis of total Na_v_1.5 protein expression normalized to GAPDH protein expression; \*\*\*\**P*<0.0001 (*One-way Anova test*); **Top.** Representative immunoblot of Na_v_1.5 expression in total protein lysates of mouse hearts. **E, Top.** Na_v_1.5 expression in the membrane fraction of Ctrl, *Scn5a*^+/-^ and *Scn5a*^+/-^ + AAV-Nter heart lysates; \**P*<0.05 (*One-way Anova test*); **Bottom**. Representative immunoblot of Na_v_1.5 expression in soluble (S) and membrane (M) fractions of heart tissues from Ctrl and *Scn5a*^+/-^ mice injected or not with AAV-Nter (MW: molecular weight). GAPDH was used as a control of soluble proteins and N-cadherin to quantify Na_v_1.5. The control mice are shown in gray, the *Scn5a*^+/-^ mice in pink and the AAV-Nter injected *Scn5a*^+/-^ mice in green.

**Table 1.**
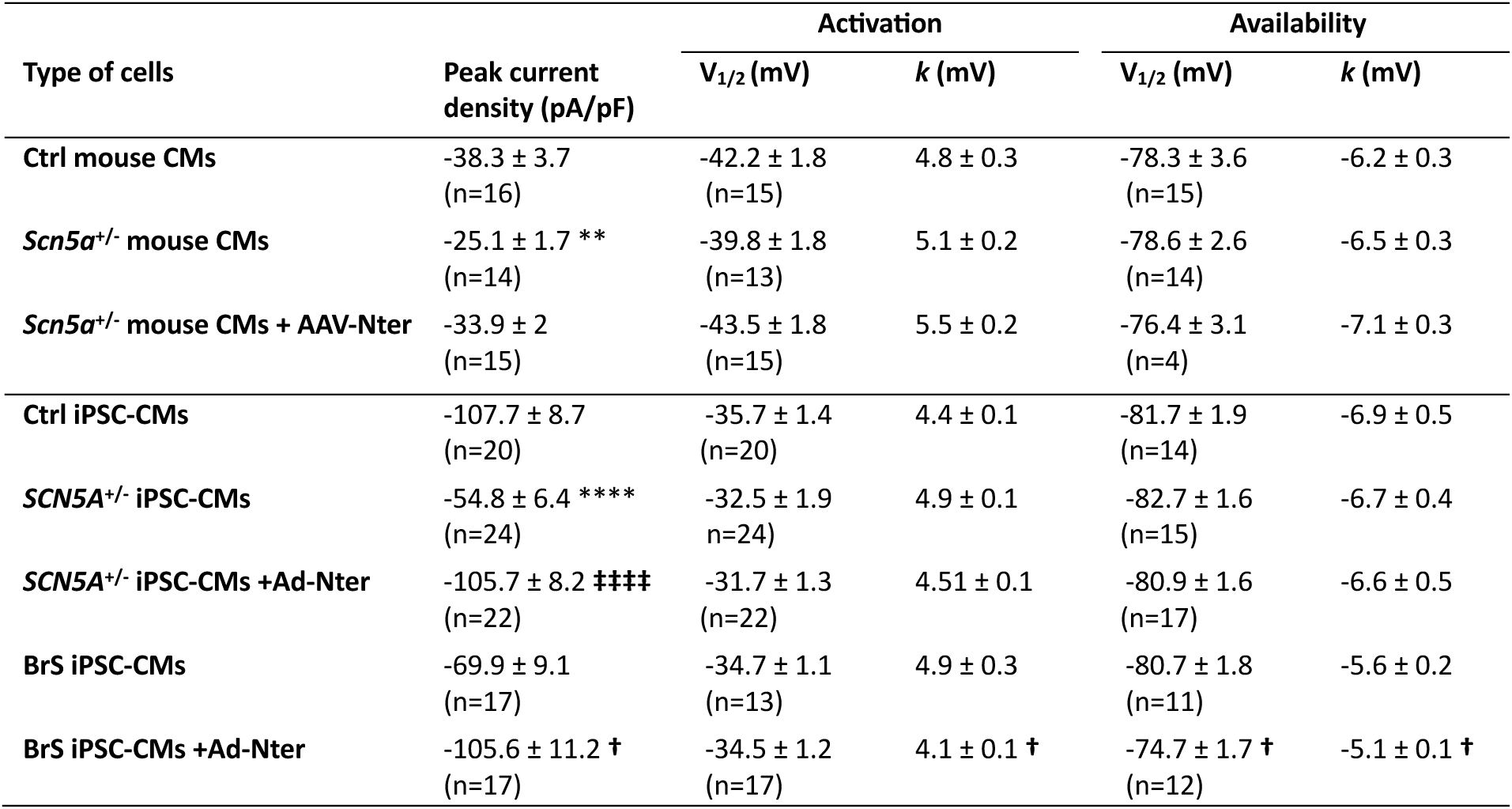
Kinetics properties of *I*_Na_ recorded in iPSC-CMs and mouse cardiomyocytes overexpressing Nter or GFP. Data are presented as means ± SEM. Peak current density is given at -20 mV for iPSC-CMs and at -30 mV for mouse cardiomyocytes. Ctrl indicates control, V_1/2_ half activation or availability value, *k* inverse slope factor, CMs cardiomyocytes and ns not significant; \**P*<0.05, ** *P*<0.01, *** *P*<0.001, **** *P*<0.0001 compared to control cells (*One-way Anova test*). ‡‡‡‡ *P*<0.0001 compared to *SCN5A*^+/-^ iPSC-CMs (*One-way Anova test*). †*P*<0.05 compared to BrS iPSC-CMs (*Student t-test*).

At the RNA and protein levels, total Scn5a transcript and Na_v_1.5 protein expression, reduced by half in *Scn5a*^+/-^ mice hearts, were unaffected by AAV-Nter treatment (Figure 1C and 1D). To investigate whether the increased *I*_Na_ recorded in AAV-Nter injected *Scn5a*^+/-^ mice was associated with a reorganization of Na_v_1.5 subcellular distribution, we quantified the cell-surface expression of Na_v_1.5 in *Scn5a*^+/-^ mice injected with AAV-Nter or with AAV-GFP. Interestingly, we observed that Na_v_1.5 surface expression was significantly increased by the Nter overexpression in comparison to GFP-injected mice (Figure 1E), suggesting that the Nter peptide promoted translocation of sodium channels to the plasma membrane. Hence, AAV-mediated Nter overexpression was efficient to normalize the specific electrophysiological phenotype associated with *Scn5a* haploinsufficiency in mice by increasing Na_v_1.5 membrane density.

### Nter protected *Scn5a*^+/-^ mice from ventricular arrhythmias

To assess whether the increase of *I*_Na_ caused by Nter overexpression was sufficient to restore ECG parameters in *Scn5a*^+/-^ mice, we recorded surface ECG. In non-treated *Scn5a*^+/-^ mice, we observed the typical ECG pattern of this BrS-like mouse model^17^, notably a prominent J-wave (Figure 2A) and a slowed cardiac conduction (Figure 2B). In Nter-treated *Scn5a*^+/-^ mice, we evidenced that Nter overexpression corrected cardiac conduction defects by normalizing the P-wave duration, the PR and QRS intervals, without modifying ECG morphology (Figure 2B and Table S3).

**Figure 2.**
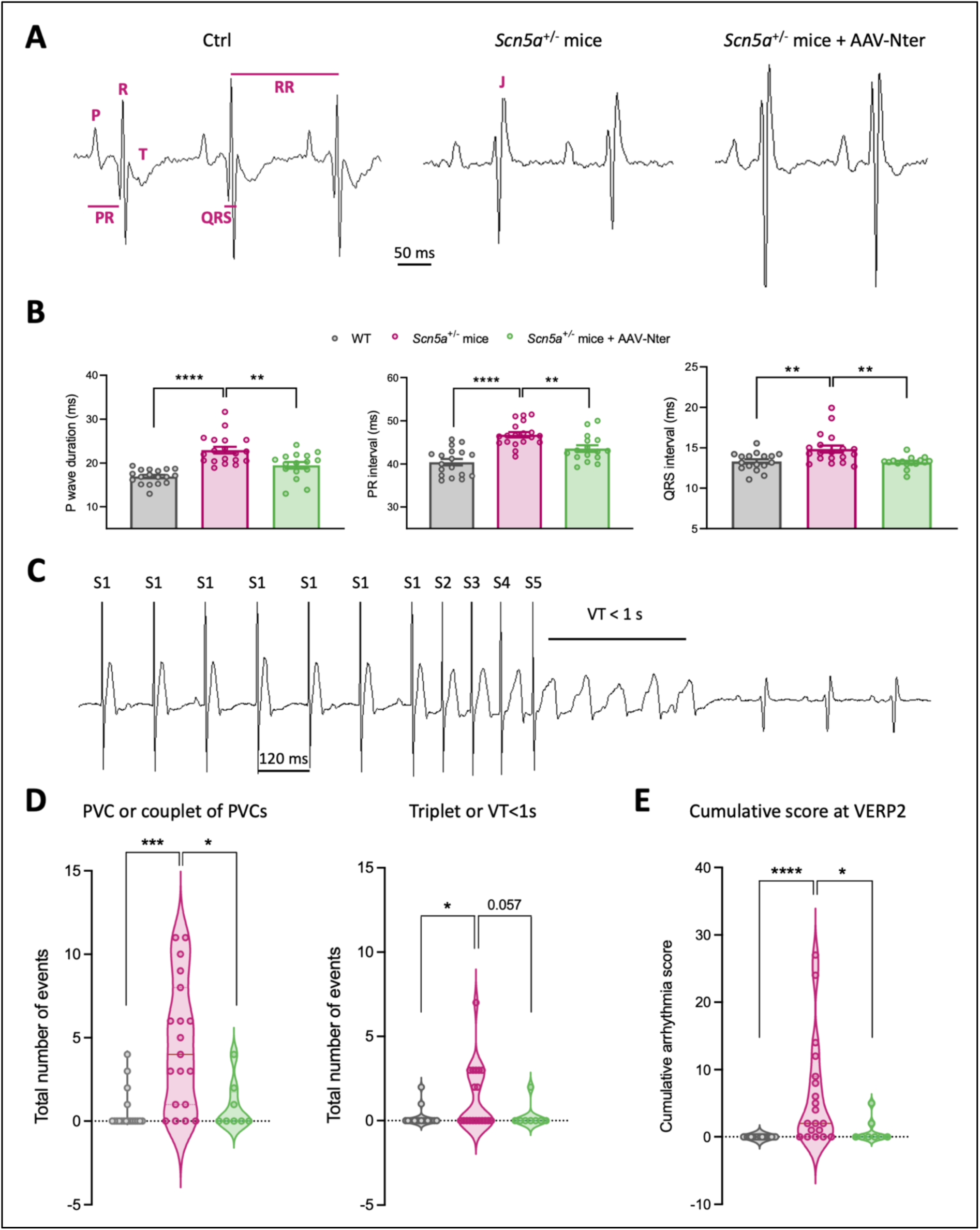
Nter overexpression prevented the occurrence of arrhythmias induced by intracardiac PES by normalizing cardiac conduction defects in *Scn5a*^+/-^ mice. **A.** Representative ECG recordings (lead I) from control (Ctrl), *Scn5a*^+/-^ mice and *Scn5a*^+/-^ mice + AAV-Nter. **B.** P-wave duration, PR and QRS intervals measured on ECG of Ctrl, *Scn5a*^+/-^ mice and *Scn5a*^+/-^ mice + AAV-Nter; \*\**P*<0.01, \*\*\*\**P*<0.0001 (*One-way Anova test*). **C.** Ventricular tachycardia (VT) trace induced by PES in a *Scn5a*^+/-^ mice. **D, Left.** Number of premature ventricular complexes (PVCs) or couplets of PVCs triggered in Ctrl (n= 18), *Scn5a*^+/-^ mice (n= 19) and *Scn5a*^+/-^ mice + AAV-Nter (n= 8); \**P*<0.05, \*\*\**P*<0.001 (*Kruskal-Wallis test*); **Right.** Number of triplets of PVCs or VTs triggered in Ctrl (n= 18), *Scn5a*^+/-^ mice (n= 19) and *Scn5a*^+/-^ mice + AAV-Nter (n= 8); \**P*<0.05 (*Kruskal-Wallis test*). **E.** Cumulative arrhythmia scores calculated in Ctrl (n= 18), *Scn5a*^+/-^ mice (n= 19) and *Scn5a*^+/-^ mice + AAV-Nter (n= 8); \**P*<0.05, \*\*\*\**P*<0.0001 (*Kruskal-Wallis test*). The control mice are shown in gray, the *Scn5a*^+/-^ mice in pink and the AAV-Nter injected *Scn5a*^+/-^ mice in green.

To further explore the Nter *in vivo* cardiac effects and its potential role as a treatment for BrS, we analyzed the *Scn5a*^+/-^-mice susceptibility to arrhythmias triggered by intracardiac programmed electrical stimulation (PES) (Figure 2C) using protocols inspired from human-clinical practice recently published^19^. The occurrence of different arrhythmic events, including premature ventricular complexes (PVC) and ventricular tachycardia (VT), was evaluated. Our results demonstrated that *Scn5a*^+/-^ mice were more likely to develop arrhythmias in response to PES than control mice and that Nter overexpression reduced the number of PVC and VT events triggered in treated mice (Figure 2D). In order to quantify the severity of arrhythmic events triggered by PES, we then calculated a cumulative arrhythmia score, as previously established^19,22^. Consistently, *Scn5a* haploinsufficiency increased the risk of arrhythmias in mice and this susceptibility was significantly limited by the Nter peptide (Figure 2E). Altogether, these results evidenced that Nter overexpression induced a peculiar protective mechanism against ventricular arrhythmias, which was correlated with a rescue of cardiac conduction, reinforcing the therapeutic potential of our strategy.

### Human model of *SCN5A* haploinsufficiency recapitulated key molecular BrS phenotype

To extrapolate our findings from the murine model to clinical applications in patients, we engineered a human cellular model of *SCN5A* haploinsufficiency by invalidating one allele of the gene using a CRISPR-Cas9 strategy in control human induced pluripotent stem cells (iPSCs)^18^. Two *SCN5A*^+/-^ iPSC lines and their isogenic control were selected for this study (Figure S3A). First, expression of pluripotency markers was verified in all clones by immunocytochemistry (Figure S3B) and RT-qPCR (Figure S3C). Cardiac differentiation efficiency was then assessed by immunocytochemistry for cardiac markers and flow cytometry. We observed that, like control iPSCs, both *SCN5A*^+/-^ iPSC lines were able to differentiate efficiently into cardiomyocytes with >85 % of troponin T-positive cells (Figure 3A) and a normal sarcomeric morphology (Figure 3B). Besides, haploinsufficiency of *SCN5A* was validated in edited iPSC-CM lines by RT-PCR and cDNA sequencing after treatment with cycloheximide, a blocker of translation (Figure S3D). Heterozygosity of both lines was confirmed by a western blot analysis showing a 2-fold reduced Na_v_1.5-protein expression in *SCN5A*^+/-^ iPSC-CMs compared with their isogenic-control lines (Figure 3C).

**Figure 3.**
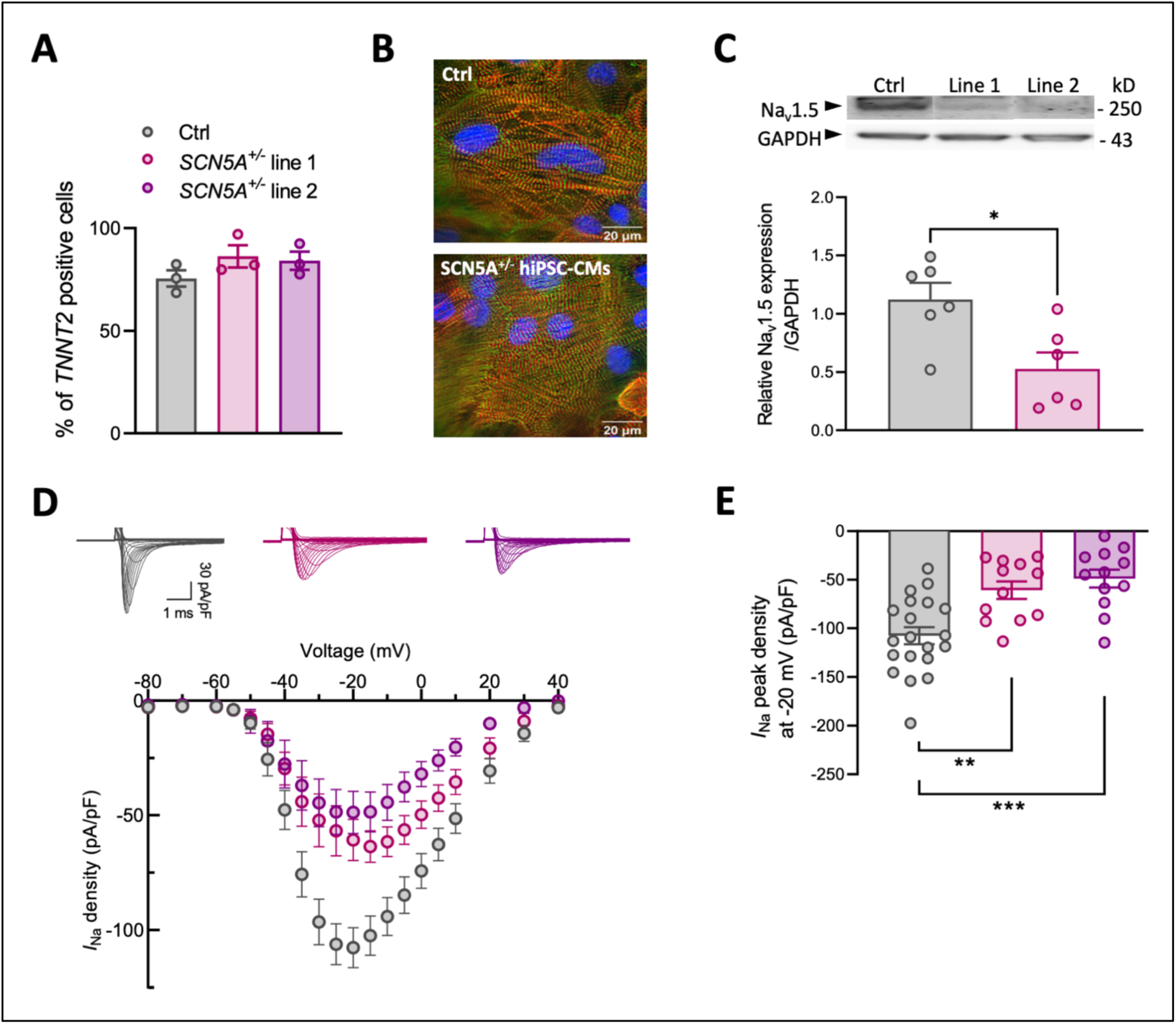
Functional characterization of cardiomyocytes differentiated from gene-edited *SCN5A*^+/-^ iPSCs. **A.** Percentage of troponin T-positive cells evaluated by flow cytometry. **B.** Immunostaining of cardiac markers α-actinin (red) and troponin T (green) in control (Ctrl) and *SCN5A*^+/-^ iPSC-CMs. **C, Top.** Representative immunoblot of Na_v_1.5 encoded by *SCN5A* from the total protein lysate at day 30 of differentiation; **Bottom.** Western-blot analysis of Na_v_1.5 expression normalized to GAPDH expression in Ctrl and *SCN5A*^+/-^ iPSC-CMs; \**P*<0.05 (*Student t-test*) **D, Top.** Representative raw traces of *I*_Na_ from Ctrl and *SCN5A*^+/-^ iPSC-CMs; **Bottom.** *I*_Na_ density-membrane potential relationships of Ctrl (n= 20) and *SCN5A*^+/-^ iPSC-CMs line 1 (n= 12) and line 2 (n= 12). **E.** Peak *I*_Na_ density measured at -20 mV in Ctrl and *SCN5A*^+/-^ iPSC-CMs; \*\**P*<0.01, \*\*\**P*<0.001 (*One-way Anova test*). All values were expressed as mean ± SEM. N=3 independent biological experiments per condition. The control iPSC-CMs are shown in gray, the *SCN5A*^+/-^ iPSC-CMs from line 1 in pink, and the *SCN5A*^+/-^ iPSC-CMs from line 2 in mauve.

At the functional level, iPSC-CMs derived from both *SCN5A*^+/-^ lines displayed a lower *I*_Na_ than control lines, mimicking a key feature of the BrS phenotype (Figure 3D and Figure 3E). As they presented a similar electrophysiological profile, the two *SCN5A*-deficient lines were considered as a single group for the rest of this study.

### Nter increased Na_v_1.5 cell-surface expression in *SCN5A*^+/-^ iPSC-CMs

To assess the effects of Nter overexpression in human *SCN5A*^+/-^ iPSC-CMs, we first analyzed the efficiency of our approach by RT-qPCR and immunostaining. As anticipated, we demonstrated a robust GFP expression in ≍95 % of iPSC-CMs transduced with the adenovirus carrying the Nter sequence (Ad-Nter), comparable to that observed with the control adenovirus Ad-GFP (Figure S4A and S4B). Similarly, the Nter-mRNA level was markedly increased in iPSC-CMs transduced with Ad-Nter (Figure S4C). Secondly, we ensured that GFP overexpression mediated by adenovirus did not alter the kinetic properties of *I*_Na_ by patch-clamp recordings (Table S4 and Figure S4D), thus validating our strategy of gene transfer in human iPSC-CMs.

At the molecular level, both *SCN5A* mRNA (Figure 4A) and Na_v_1.5 total proteins (Figure 4B) were reduced by half in *SCN5A*^+/-^ iPSC-CMs and Nter overexpression did not modify their expression levels, indicating that Nter was not involved in modulating Na_v_1.5 transcription or translation mechanisms. In contrast, immunocytochemistry analysis of Na_v_1.5 cell-distribution emphasized a restoration of Na_v_1.5-membrane expression by overexpression of the Nter peptide in *SCN5A*^+/-^ iPSC-CMs (Figure 4C). Therefore, our results obtained in iPSC-CMs confirmed that Nter peptide promoted Na_v_1.5 expression at the plasma membrane, as observed in the *Scn5a*^+/-^ murine model.

**Figure 4.**
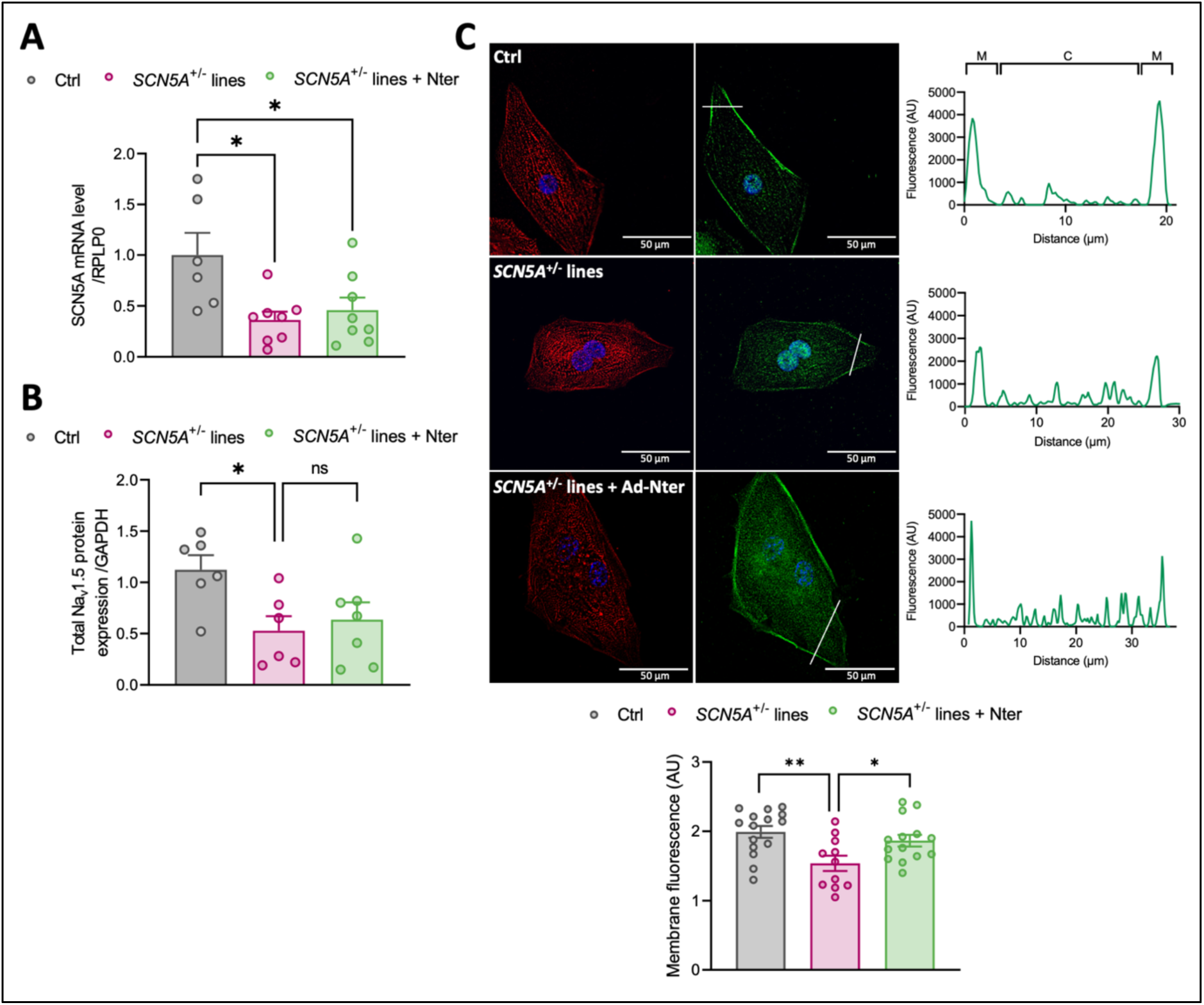
Na_v_1.5 membrane expression was increased by Nter overexpression in *SCN5A*^+/-^ iPSC-CMs. **A.** *SCN5A* mRNA expression level in control (Ctrl) and *SCN5A*^+/-^ iPSC-CMs transduced with Ad-GFP or Ad-Nter. Cts were normalized to h*RPLP0* and the ratio *vs.* Ctrl (2^-ΔΔCt^) was then calculated; \**P*<0.05 (*One-way Anova test*) **B.** Western-blot analysis of Na_v_1.5 expression normalized to GAPDH expression in Ctrl and *SCN5A*^+/-^ iPSC-CMs transduced with Ad-GFP or Ad-Nter; \*\**P*<0.01 (*One-way Anova test*). **C, Left.** Representative confocal images for immunostainings of α-actinin (red) and Na_v_1.5 (green) in Ctrl and *SCN5A*^+/-^ iPSC-CMs transduced with Ad-GFP or Ad-Nter; **Right.** Fluorescence measurement of Na_v_1.5 staining using ImageJ software; **Bottom.** Quantification of Na_v_1.5 expression at the plasma membrane in Ctrl and *SCN5A*^+/-^ iPSC-CMs transduced with Ad-GFP or Ad-Nter; \**P*<0.05, \*\**P*<0.01 (*One-way Anova test*). All values were expressed as mean ± SEM. N = 3 independent biological experiments per condition. The control iPSC-CMs are shown in gray, the *SCN5A*^+/-^ iPSC-CMs + Ad-GFP in pink and the *SCN5A*^+/-^ iPSC-CMs + Ad-Nter in green.

### Nter restored electrophysiological properties of *SCN5A*^+/-^ iPSC-CMs

To assess the functional effects of Nter overexpression in *SCN5A*^+/-^ iPSC-CMs, we then recorded *I*_Na_ and APs in single cells. We evidenced that Nter normalized *I*_Na_ density (Figure 5A and 5B), without affecting the current biophysical parameters (Table 1). In line with these results, we observed that Nter peptide restored dV/dt_max_ to control levels in *SCN5A*^+/-^ iPSC-CMs (Figure 5C). In contrast to APs recorded in *Scn5a*^+/-^ mice, APs recorded in *SCN5A*^+/-^ iPSC-CMs were prolonged at 10, 30, 50 and 90 % of repolarization, compared with control cells. Notably, Nter overexpression partially restored APDs to control levels, since APD90 was significantly shortened (Figure 5D). Besides, resting membrane potential, AP amplitude and overshoot were not modified by *SCN5A* haploinsufficiency and Nter overexpression (Figure S5A, S5B and S5C). Importantly, Nter peptide suppressed early after depolarizations (EADs) recorded in non-treated *SCN5A*^+/-^ iPSC-CMs (Figure 5E). Altogether, our results showed that Nter peptide normalized the electrophysiological phenotype of *SCN5A*-haploinsufficient cells. Finally, in order to challenge our gene-therapy approach in a human pathophysiological context, we overexpressed Nter in iPSC-CMs derived from a BrS patient (BrS iPSC-CMs) carrying a haploinsufficient mutation in *SCN5A*^23^. Again, we observed that Nter overexpression significantly increased *I*_Na_ density in BrS iPSC-CMs (Figure 6A and 6B), while *I*_Na_ activation was unaffected, and its availability was slightly shifted towards more positive potentials (Table 1). Interestingly, Nter overexpression abolished EAD occurrence observed in 40% of spontaneous APs recorded in BrS iPSC-CMs (Figure 6C). Overall, our *in vitro* results suggest that overexpression of the Nter peptide promotes Na_v_1.5 expression at the cell surface sufficiently to reverse the BrS phenotype associated with *SCN5A* haploinsufficiency.

**Figure 5.**
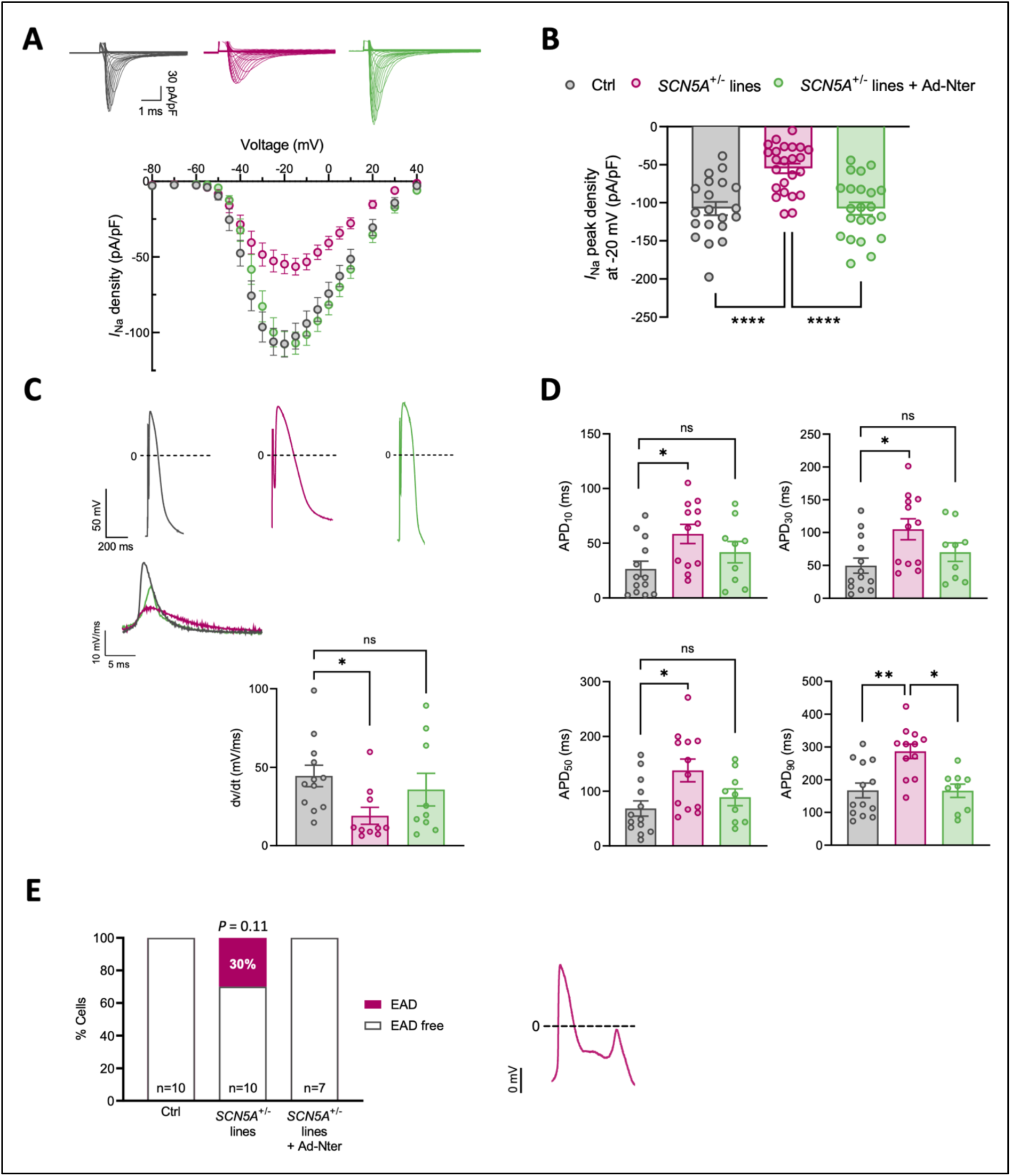
Adenovirus-mediated Nter overexpression normalized electrophysiological properties in *SCN5A*^+/-^ iPSC-CMs. **A, Top.** Representative raw traces of *I*_Na_ from Ctrl and *SCN5A*^+/-^ iPSC-CMs transduced with Ad-GFP or Ad-Nter; **Bottom.** Current density-membrane potential relationships of Ctrl (n= 20) and *SCN5A*^+/-^ iPSC-CMs transduced with Ad-GFP (n= 24) or Ad-Nter (n= 22). **B.** Peak *I*_Na_ density measured at -20 mV in Ctrl and *SCN5A*^+/-^ iPSC-CMs transduced with Ad-GFP or Ad-Nter; \*\*\*\**P*<0.0001 (*One-way Anova test*). **C, Top**. Representative AP and maximum upstroke velocity (dV/dt_max_) traces from Ctrl and *SCN5A*^+/-^ iPSC-CMs transduced with Ad-GFP or Ad-Nter; **Bottom.** dV/dt_max_ in Ctrl and *SCN5A*^+/-^ iPSC-CMs transduced with Ad-GFP or Ad-Nter; \**P*<0.05 (*Kruskal-Wallis test*). **D**. AP durations (APDs) at 10 (APD_10_), 30 (APD_30_), 50 (APD_50_) and 90 % (APD_90_) of repolarization recorded in Ctrl and *SCN5A*^+/-^ iPSC-CMs transduced with Ad-GFP or Ad-Nter; \**P*<0.05, \*\**P*<0,01 (*Kruskal-Wallis test*). **E, Left**. Percentage of EADs in Ctrl and *SCN5A*^+/-^ iPSC-CMs transduced with Ad-GFP or Ad-Nter (*Fisher one-tail test*). All values were expressed as mean ± SEM.; **Right.** Representative early-after depolarization (EAD) during spontaneous AP recordings. N = 3 independent biological experiments per condition. The control iPSC-CMs are shown in gray, the *SCN5A*^+/-^ iPSC-CMs + Ad-GFP in pink and the *SCN5A*^+/-^ iPSC-CMs + Ad-Nter in green.

**Figure 6.**
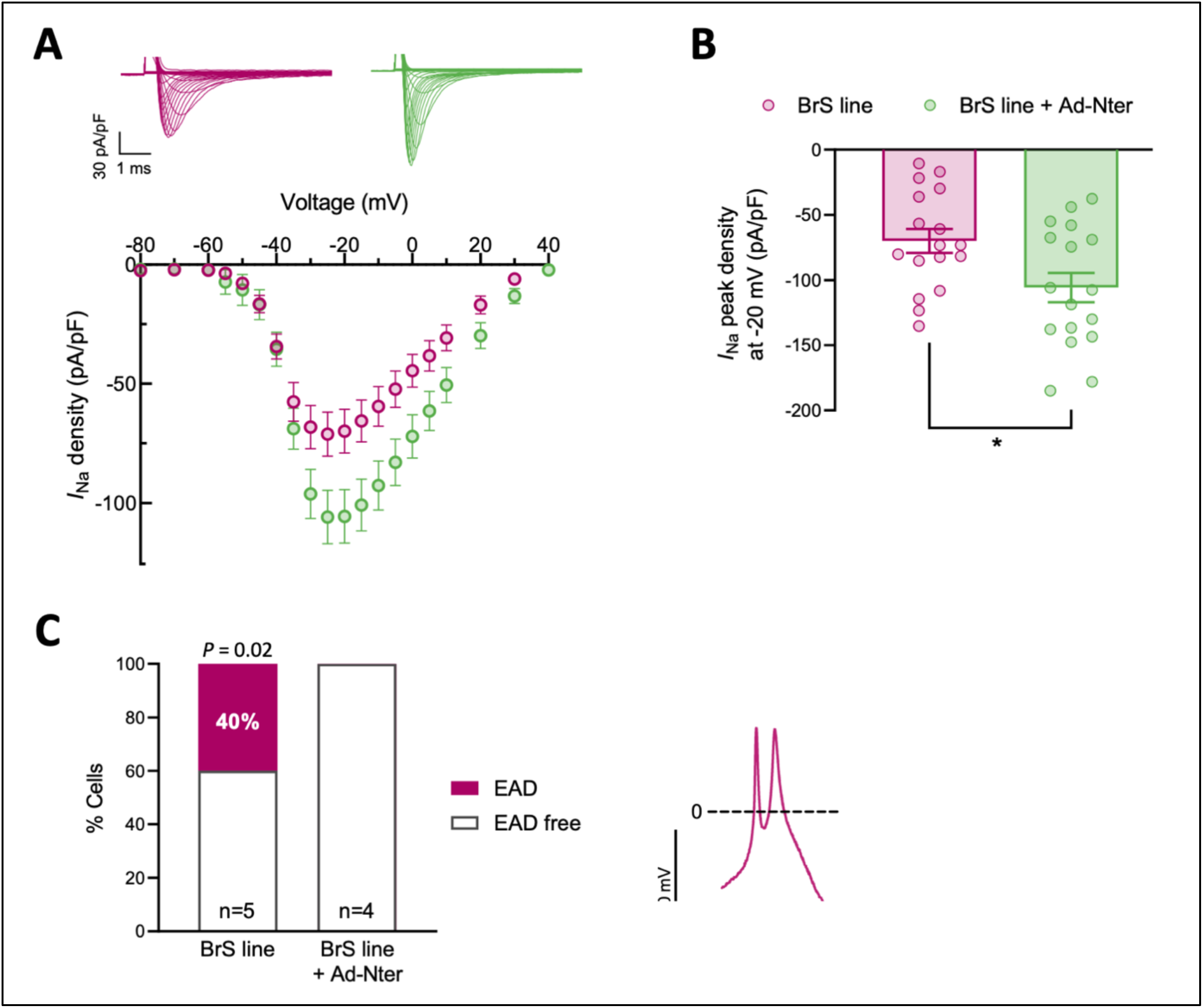
Adenovirus-mediated Nter overexpression normalized electrophysiological properties in iPSC-CMs from a BrS patient. **A, Top**. Representative raw traces of *I*_Na_ from BrS iPSC-CMs transduced with Ad-GFP or Ad-Nter; Bottom. Normalized current density-membrane potential relationships of BrS iPSC-CMs transduced with Ad-GFP (n= 17) or Ad-Nter (n= 17). **B.** Peak *I*_Na_ density measured at -20 mV in BrS iPSC-CMs transduced with Ad-GFP or Ad-Nter; \**P*<0.05 (*Student t test*). **C, Left**. Percentage of EADs in BrS iPSC-CMs transduced with Ad-GFP or Ad-Nter (*Fisher one-tail test*); **Right**. Representative early-after depolarization (EAD) during spontaneous AP recordings. All values were expressed as mean ± SEM. N = 3 independent biological experiments per condition. The BrS iPSC-CMs + Ad-GFP are shown in pink and the BrS iPSC-CMs + Ad-Nter in green.

## DISCUSSION

Our study highlights a novel gene therapy strategy to reverse the BrS phenotype associated with *SCN5A* haploinsufficiency. We developed a human cellular model representative of 40-50% of BrS mutations identified in *SCN5A* and combined its study with the heterozygous *Scn5*a knockout mouse, a BrS-like murine model. In both models, we have demonstrated that the overexpression of the N-terminal fragment of Na_v_1.5 (Nter) can successfully reverse BrS-associated cardiac cellular and *in vivo* electrophysiological abnormalities. AAV-Nter facilitated cell-surface expression of full-length Na_v_1.5 native channels and corrected electrophysiological defects through normalization of *I*_Na_ density and AP parameters. Most importantly, our findings established that arrhythmias can be successfully prevented *in vivo* and *in vitro* in models of *SCN5A* haploinsufficiency by the Nter peptide.

Development of drug therapy to prevent life-threatening arrhythmias is a very complex field in pharmacology and, to date, no antiarrhythmic drugs are effective in preventing SCD except for β-blockers, whose action is not specifically targeted to the arrhythmogenic substrate. The use of ICDs is an effective therapy for reducing SCD and these devices play a key role in preventing premature death in patients with inherited arrhythmias^10^. However, due to a fairly high percentage of adverse effects, especially in young patients, ICDs are targeted to patients identified as having a high risk of developing life-threatening arrhythmias and SCD. Nevertheless, the risk of SCD remains challenging to predict, emphasizing the necessity for novel treatments to reduce morbidity and mortality in these young, otherwise healthy patients with channelopathies. The development of gene therapy approaches may offer a potential solution, as proposed over the past decade^24–26^. The best known example of effective gene therapy for channelopathies is a replacement strategy based on the *CASQ2* gene overexpression using viral vectors in animal and human cellular models of catecholaminergic polymorphic ventricular tachycardia^25,27^. In these studies, transduction of mice at birth completely prevented phenotypic features of the disease, and mice injected as adults showed complete regression of disease manifestations, including life-threatening ventricular arrhythmias, even one year after administration of the viral construct^27^. This ground-breaking research has paved the way for further studies dedicated to gene therapy of channelopathies. Since 1998, when the first mutations in *SCN5A* were identified^28^, very few therapeutic strategies have been proposed to directly target the dysfunctional sodium channel activity. Although replacement is the most straightforward gene therapy approach, it is difficult to apply to *SCN5A* because the gene size (>6 kb) exceeds the packaging capacity (≍5 kb) of AAVs, which are the most suitable vectors for human gene therapy due to their superior safety profile^29^. *In vivo* transfer of the full-length *SCN5A* gene using a trans-splicing and viral DNA recombination strategy is feasible^14^, but its potential side effects, in particular unwanted gene products, make its translation to clinical applications difficult. In a recent study, Yu *et al.* proposed the first gene therapy for the treatment of BrS using AAV9-mediated expression of MOG1, a small chaperone protein of Na_v_1.5. The authors were able to rescue *I*_Na_ in cellular and animal models carrying the heterozygous *SCN5A* mutation G1743R, which is responsible for a defect in Na_v_1.5 trafficking^30^. It should be noted that *MOG1* gene expression was driven by the cytomegalovirus promoter, which could induce non-cardiac-specific expression of the ubiquitous MOG1 protein, leading to potential adverse side effects, despite the use of AAV serotype 9, which is characterized by a strong tropism for cardiac tissue in mice^31^. In the present study, we ensured cardiac-specific expression of Nter, an exogenous peptide known to increase *I*_Na_^15,16^, by using the cTnT promoter in conjunction with AAV serotype 9 and confirmed the efficacy and safety of this cardiac gene delivery vector, which avoided Nter expression in non-cardiac tissues, particularly the brain. As a result, we emphasized that Nter acts as a molecular driver of Na_v_1.5 protein expression at the cell surface, leading to increased *I*_Na_ density and reversion of the BrS-like phenotype associated with *SCN5A* haploinsufficiency in mouse and human cellular models. The same phenomenon of transcomplementation has been previously investigated in the context of cystic fibrosis, with publications demonstrating the use of a CFTR chloride channel fragment to enhance the expression of mutated channels and partially restore their function^32,33^.

One limitation of our approach is that haploinsufficiency does not fully account for the spectrum of *SCN5A* mutations identified in BrS patients. Nevertheless, this strategy presents a significant opportunity for treating 40-50% of patients using a single gene-therapy vector. Another advantage of our approach is that Nter overexpression favors cell-surface expression of native Na_v_1.5 proteins without increasing *SCN5A* transcription or sodium channel translation. This prevents overproduction of Na_v_1.5, which could otherwise be arrhythmogenic. Furthermore, it could be extended to cardiac diseases in which conduction disorders are not directly related to haploinsufficiency of *SCN5A* but where the Na^+^-channel expression is altered, such as arrhythmogenic cardiomyopathies, atrial fibrillation or heart failure^34–36^.

The main finding of this study is that Nter overexpression markedly decreases the incidence of arrhythmic events, which mirror the pivotal characteristics of BrS, in both murine and cellular models. Although arrhythmias are unlikely to occur spontaneously in mice^37^, we chose not to use aggressive PES protocols, as the results may be nonspecific and yield false positives^38^. Instead, we designed protocols based on human clinical practice. These protocols bring extrastimuli close to the refractory period of the stimulated ventricular area, which favors unidirectional propagation block serving as re-entry phenomenon substrate. Moreover, such PES protocols are employed to induce a re-entry phenomenon, which represents the most prevalent arrhythmic mechanism sustaining ventricular fibrillation^39^. More specifically, a positive PES has been shown to be associated with a higher risk of ventricular arrhythmia occurrence in Brugada patients^40,41^.

In this study, we recorded prolonged APDs associated with EADs in *SCN5A*^+/-^ iPSC-CMs. EADs are related to abnormal depolarization of the cardiac tissue occurring during phase 2 or 3 of the cardiac AP. They develop in the setting of AP prolongation and, when the sodium current reactivates, can trigger premature ventricular beats and ventricular tachycardias. These triggered activities are not typically associated with BrS-related arrhythmias in humans, or in *Scn5a*^+/-^ mice. However, these phenotypic characteristics have been observed in several iPSC-CMs models of BrS^42–44^, a discrepancy that could be attributed to the characteristic immaturity of this cellular model, but also to the nature of the sample (isolated cells), underlying an influence of the experimental conditions and model^45^. It is noteworthy that these EADs were observed in a context of APD prolongation of spontaneous APs recorded in *SCN5A*^+/-^ iPSC-CMs. In contrast, haploinsufficiency did not prolong murine APD at 5% and 50% of repolarization and even slightly shortened it at 90% of repolarization, and no EADs were observed, consistent with a BrS-like phenotype. This observation could be the result of a different influence of the repolarizing currents I_to_ and I_K1_ on APD in mice *versus* iPS-CMs. Indeed, another limitation of the iPSC-CM cellular model is the absence of the hyperpolarizing *I*_K1_ current, which prevents the recruitment of all the channels classically involved in the AP. However, the absence of a difference in overshoot and AP amplitude between groups of iPSC-CMs provided evidence that the same percentage of ion currents were recruited in all groups. Moreover, the pathophysiological mechanism of BrS originates in the human right ventricular outflow tract, but current cardiac differentiation protocols of iPSCs do not allow chamber (left or right ventricle/atria) and layer (sub-endocardium, midmyocardium, sub-epicardium) specifications.

Despite their limitations in recapitulating the intricate structural and functional aspects of the mature heart *in vitro*, human iPSC-CMs effectively recapitulate the phenotype of various inherited cardiac arrhythmias, making them a widely used research tool^46^. Finally, it was of interest that the EADs observed in spontaneous APs recorded *in vitro* were abolished by Nter overexpression. Furthermore, it is important to note that the results were fully replicated in a cell line derived from a Brugada patient. The results presented herein provide compelling evidence that restoration of Na_v_1.5 expression at the plasma membrane through Nter overexpression is an effective strategy for alleviating the arrhythmogenic substrate and preventing arrhythmias in *SCN5A*-haploinsufficient cells. Further studies are required to ascertain the underlying mechanisms responsible for the observed effects of the Nter peptide on Na_v_1.5 subcellular expression and to elucidate its role, either as a chaperone^15^ or as a decoy, in facilitating the translocation of an existing pool of functional channels to the cell surface. In this regard, the recent study by Witmer *et al*. is of particular interest for its findings regarding the existence of a peptide of Na_v_1.5 endogenously expressed in humans, but not in rodents^47^. The peptide is composed of the N-terminus of the channel and a C-terminus derived from an intronic sequence. Furthermore, the study indicates that this previously unidentified short protein exerts an influence on mitochondrial physiology, but does not affect *I*_Na_ density or Na_v_1.5 expression, in contrast to the Nter.

In conclusion, we developed a gene therapy based on AAV-mediated overexpression of the cardiac sodium channel N-terminal region as a potential treatment for *SCN5A* haploinsufficiency. AAV-Nter vectors promoted cell surface expression of Na_v_1.5, increased sodium current density and suppressed EADs in *SCN5A*^+/-^ iPSC-CMs and in iPSC-CMs from a Brugada patient. Furthermore, AAV-Nter gene therapy was demonstrated to increase cell surface Na_v_1.5 expression, enhance *I*_Na_ density, normalize ECG parameters and protect *Scn5a*^+/-^ mice from triggered arrhythmias. Our study demonstrates the potential benefits of increasing the cardiac sodium channel Na_v_1.5 at the cell surface of cardiomyocytes to reverse the electrophysiological disease phenotype associated with *SCN5A* haploinsufficiency. A comparable approach could be applied to other cardiac diseases associated with a deficiency of the sodium current.

## ACKNOWLEDGEMENTS

The authors are grateful to Dr. A. Coulombe (Paris, France) and to Dr. E. Balse (Paris, France) for their expertise in cardiac electrophysiology and to Dr. E. Villard (Paris, France) and L. Duboscq-Bidot (Paris, France) for their advice on CRISPR/Cas9 editing. The authors also thank the iPS-ICAN facility for technical help, the Viral Vector Core of Nantes University (Nantes, France) for AAV and adenovirus production, Dr. B. French (Virginia University, USA) for the gift of plasmid pAcTnT-eGFP and Dr. A. Grace for the authorization to work with the *Scn5a*^+/-^ mouse line. The authors are also grateful to Doriane Joubert-Foret and Yannick Martinez from UMS-28 (NAC) for animal care.

## SUPPLEMENTAL INFORMATION

Expanded Methods

Supplemental Tables: Table S1-S6

Supplemental Figures: Figure S1-S5

