## Supplemental data for "Gene Therapy with the N-terminal Fragment of Na_v_1.5 for Cardiac Channelopathies: A Novel Transcomplementation Mechanism Potentiating the Cardiac Sodium Current"

**SUPPLEMENTAL MATERIAL**

Expanded Methods

Supplemental Tables: Table S1-S6

Supplemental Figures: Figure S1-S5

**EXPENDED MATERIALS AND METHODS**

**Viral vectors**

AAV9 and adenoviral vectors were produced by the Viral Vector Core of Nantes University (*France*). The AAV vector pAcTnT-eGFP, used as control (AAV-GFP), has been described previously[14]. The AAV-Nter carries the first 132 amino acids of the human sequence hH1a (Nter) of $Na_V1.5$ (RefSeq accession number NM_000335.4) separated from the eGFP reporter gene by an internal ribosome entry site (IRES). The Nter sequence is under the control of the cardiac-specific chicken troponin T2 promoter. In adenovirus Ad-Nter, the human Nter coding sequence fused to a tag flag was cloned upstream of an IRES and the eGFP reporter gene. This vector allows Nter-flag and eGFP expression driven by the Cytomegalovirus (CMV) promoter. Adenovirus expressing eGFP alone was used as control (Ad-eGFP).

**BrS-patient derived iPS-CMs**

*SCN5A*$^{+/A655G\text{-}fsX16}$ iPSCs presented in this paper were reprogrammed from peripheral blood mononuclear cells of a Brugada patient and have been previously characterized[23]. It should be noted that we chose to use this BrS patient iPS line despite the absence of an isogenic control line because this study focuses on the potentiating effects of Nter-peptide overexpression on $I_{Na}$ and we considered that comparison of treated cells with a healthy control was not essential.

**Generation of *SCN5A*-deficient induced pluripotent stem cells**

By CRISPR/Cas9 genome editing of a control line, we engineered iPSCs with a heterozygous frameshift mutation in exon 2 of *SCN5A*, leading to a premature stop codon and *SCN5A*-haploinsufficiency as we previously published[18]. A PCR-based method was developed using a forward primer 5′-GAATCAGGCCCATTGTCTGT-3′ and a reverse primer 5′-GTGACTCATTTCCCCAGAGC-3′ for targeted mutation identification by Sanger sequencing (*Eurofins Genomics*). The present study was conducted on two edited *SCN5A*$^{+/-}$ iPSC clones, carrying the mutation p.A15S-fsX81 for clone 1 and the mutation p.F16V-fsX72 for clone 2, both of which are absent from databases listing variants in *SCN5A* (Fig. S2). The genomic integrity, maintenance of pluripotency, and differentiation potential of the *SCN5A*$^{+/-}$ clones have been described elsewhere[18].

**Echocardiography**

Eight weeks after injection, male and female mice were anesthetized with isoflurane for echocardiography. Parameters were evaluated by the Vivid7 PRO echocardiography Doppler device (*GE Medical System*) with an ultrasound-emitting probe (frequency between 9 and 14

MHz). The time-motion mode recording of the left ventricle (LV) in the parasternal long-axis view, allowed various parameters measurements: diastolic and systolic posterior wall thicknesses of LV (dPWT-LV and sPWT-LV), relative wall thickness (RWT), diastolic and systolic LV diameter (dLVD and sLVD), stroke volume (SV), ejection fraction (EF) and heart rate (HR). At least 3 sets of measurements were made on 3 different cardiac cycles for each mouse. The shortening fraction (SF) was calculated with the following formula: [(LVEDD – LVESD)/LVEDD] × 100.

**Programmed electrical stimulation**

After surface ECG recording, etomidate-anesthetized mice (*12 mg/kg*) were placed on a heating pad at 37°C. After a subcutaneous injection of Xylocaine (*93 mg/kg*) around the surgical area, a small chest incision was made to expose and isolate the right jugular vein. A 1.1 octopolar catheter was inserted into the jugular vein to the right ventricular apex. Once the catheter was positioned, voltage pulses were applied by an external stimulator (*Multichannel Systems*) under the control of the MCStimulus II software (*Multichannel Systems*). A first protocol was performed by decreasing voltage amplitude from 2500 to 150 mV until the ventricular capture was lost to identify the excitation threshold. The ventricular refractory period was then determined by delivering a train of 20 stimuli (S1) at a pacing cycle length corresponding to 70 % of the animal RR interval, followed by a single premature extrastimulus (S2) with an S1-S2 cycle length shortened by 2-ms steps until the ventricular capture was lost. All protocols used in this study were inspired by human clinical practice and were based on the repetition of 2 to 4 extrastimuli, *i.e.* S2 to S5, following the repetition of 20 S1 stimuli to assess the susceptibility of *Scn5a*$^{+/-}$ mice to ventricular arrythmias compared with control mice, as recently published[19]. Triggered premature ventricular complexes (PVCs) were then quantified using the ecgAUTO software (*emka TECHNOLOGIES*).

In order to quantify the severity of arrhythmic events triggered by PES, we have calculated a score for each stimulation protocol, as established by Clasen *et al.* in 2018[22]. This score assigns 1 point for an isolated PVC, 3 points for a couplet, 4 points for a VT of 3 consecutive PVCs, 5 points for a VT of at least 5 PVCs and 6 points for a VT lasting more than 1 sec. Since none of the mice in our study triggered VT approaching a duration of 1 sec, we adapted the Clasen's score by assigning 4 points for VTs of 3 consecutive PVCs and 5 points for VTs of more than 4 complexes. For each mouse, a cumulative arrhythmia score was calculated by adding the score calculated for each stimulation protocol (S2 to S5)[19].

**Immunochemistry**

Immunostaining was performed on mouse ventricle cryosections or iPSC-CMs fixed with paraformaldehyde 4% for 10 minutes at room temperature and washed 3 times with PBS. Cells were then permeabilized in blocking solution (PBS, 2 % Bovine Serum Albumin for iPSC or 5 % for mouse ventricles, 1 % Triton) for 30 min and stained with primary antibodies overnight at 4°C. Detection was performed after 3 PBS washes and 1 hour of incubation with secondary antibodies and DAPI for nuclear staining at room temperature. Images were acquired with the 3D high-resolution microscope Deltavision (*GE healthcare*) and analyzed with ImageJ software. All antibodies used for immunostaining are listed in Table S5.

**RT-qPCR analysis**

Total mRNAs were extracted from murine heart tissues following QIAzol lysis protocol (*Qiagen*) and from iPSC-CMs using the RNeasy® mini kit (*Qiagen*). After DNase treatment (*Promega*), RNA was reverse transcribed using the Superscript II First Strand Synthesis System (*Invitrogen*) according to the manufacturer's protocols. Gene expression was assessed by qPCR using SYBR Green PCR Master Mix (*Applied Biosystems*) with the Quantstudio™ 3 Real-Time PCR instrument (*Applied Biosystems*) and calculated using the ΔΔCt method[48]. Primers used are listed in Table S6.

**Preparation of total protein and membrane protein extracts**

For total protein extraction, murine heart tissues or iPSC-CMs were homogenized in lysis buffer (50 mM Tris-HCL, 150 mM NaCl, 2 mM EDTA, 1 % Triton) supplemented with protease inhibitor cocktail (*Roche*) and incubated at 4°C for 1 hour with agitation. The lysates were centrifuged at 13000 G and at 4°C for 30 min. The supernatant was collected and total proteins were quantified using the Pierce™ BCA Protein Assay kit (*Pierce*).

Membrane protein extraction from murine heart tissues was performed using the Proteoextract® Native Membrane Protein Extraction kit (*Sigma-Aldrich*) in accordance with the manufacturer's instructions. Briefly, murine hearts were gently homogenized in the kit Extraction Buffer I with a pre-cooled Dounce glass homogenizer (*Life Sciences*). An initial centrifugation was realized to separate soluble proteins. The pellet, corresponding to a majority of cell membranes, was resuspended in Extraction Buffer II, followed by a second centrifugation to isolate a majority of cell-surface proteins.

**Western blot analysis**

Proteins were resolved by SDS-PAGE electrophoresis on NuPAGE™ 7 % Tris-acetate gels or NuPAGE™ 4-12 % Bis-Tris gels (*Invitrogen*) and transferred to nitrocellulose membranes (*Biorad*) before incubation with primary antibodies overnight at 4°C. After 3 washes with PBS-0,1 % Tween, the membranes were incubated with secondary antibodies for 1 hour at room temperature. For murine proteins, infrared dye-conjugated secondary antibodies were used, and signals were detected with the Li-COR Odyssey Infrared Imaging system (*Licor*). For iPSC-CM proteins, HRP-conjugated secondary antibodies were used, and the signal was detected with the ImageQuant™ LAS-4000 instrument (*GE Healthcare*). Western blotting images were analyzed with ImageJ software. All antibodies used for western blot are listed in Table S5.

**SUPPLEMENTAL TABLES**

| | HR (bpm) | dPWT-LV (mm) | sPWT-LV (mm) | dLVD (mm) | sLVD (mm) | SV (ml) | RWT | EF (%) | SF (%) |
|---|---|---|---|---|---|---|---|---|---|
| **Non-injected WT mice (n = 7)** | 630 ± 7 | 0.63 ± 0.04 | 1.1 ± 0.05 | 3.4 ± 0.1 | 1.8 ± 0.07 | 0.086 ± 0.007 | 0.37 ± 0.02 | 84 ± 0.6 | 47 ± 0.7 |
| **WT mice + AAV-GFP (n = 7)** | 629 ± 7 | 0.6 ± 0.03 | 1.1 ± 0.06 | 3.4 ± 0.08 | 1.8 ± 0.06 | 0.083 ± 0.005 | 0.37 ± 0.01 | 84 ± 0.5 | 47 ± 0.6 |
| **Non-injected *Scn5a*$^{+/-}$ mice (n = 9)** | 618 ± 5 | 0.62 ± 0.02 | 1.0 ± 0.02 | 3.2 ± 0.07 | 1.7 ± 0.04 | 0.073 ± 0.004 | 0.39 ± 0.01 | 84 ± 0.3 | 47 ± 0.3 |
| ***Scn5a*$^{+/-}$ mice + AAV-GFP (n = 10)** | 622 ± 5 | 0.63 ± 0.03 | 1.1 ± 0.02 | 3.3 ± 0.08 | 1.7 ± 0.05 | 0.079 ± 0.005 | 0.38 ± 0.01 | 85 ± 0.4 | 48 ± 0.4 |
| **Ctrl WT mice (n = 14)** | 630 ± 5 | 0.61 ± 0.02 | 1.1 ± 0.04 | 3.4 ± 0.07 | 1.8 ± 0.04 | 0.084 ± 0.004 | 0.37 ± 0.01 | 84 ± 0.4 | 47 ± 0.4 |
| **Ctrl *Scn5a*$^{+/-}$ mice (n = 19)** | 620 ± 4 | 0.63 ± 0.02 | 1.1 ± 0.02 | 3.2 ± 0.05 | 1.7 ± 0.03 | 0.076 ± 0.003 | 0.39 ± 0.006 | 84 ± 0.3 | 47 ± 0.3 |

**Table S1. Echocardiographic parameters of WT and *Scn5a*$^{+/-}$ mice injected or not with AAV-GFP.** Data are presented as mean ± SEM. Control (Ctrl) mice include non-injected and GFP-injected mice. WT indicates wild type, HR Heart Rate, dPWT-LV diastolic Posterior Wall Thickness-Left Ventricle, sPWT-LV systolic Posterior Wall Thickness-Left Ventricle, dLVD diastolic Left Ventricle Diameter, sLVD systolic Left Ventricle Diameter, SV Stroke Volume, RWT relative wall thickness, EF Ejection Fraction and SF Shortening Fraction. Since no significant differences were observed between AAV-GFP-injected and non-injected mice, we considered AAV-GFP-injected mice as the control group in the rest of this study (*Mann Whitney test*).

| | HR (bpm) | dPWT-LV (mm) | sPWT-LV (mm) | dLVD (mm) | sLVD (mm) | SV (ml) | RWT | EF (%) | SF (%) |
|---|---|---|---|---|---|---|---|---|---|
| **Ctrl WT mice (n = 14)** | 630 ± 5 | 0.61 ± 0.02 | 1.1 ± 0.04 | 3.4 ± 0.06 | 1.8 ± 0.04 | 0.084 ± 0.004 | 0.37 ± 0.01 | 84 ± 0.4 | 47 ± 0.4 |
| **WT mice + AAV-Nter (n = 15)** | 629± 4 | 0.66 ± 0.02 | 1.1 ± 0.02 | 3.3 ± 0.05 | 1.8 ± 0.03 | 0.081 ± 0.004 | 0.39 ± 0.01 | 84 ± 0.5 | 47 ± 0.6 |
| **Ctrl *Scn5a*$^{+/-}$ mice (n = 19)** | 620 ± 4 | 0.63 ± 0.02 | 1.1 ± 0.02 | 3.2 ± 0.05 | 1.7 ± 0.03 | 0.076 ± 0.003 | 0.39 ± 0.01 | 84 ± 0.3 | 47 ± 0.3 |
| ***Scn5a*$^{+/-}$ mice + AAV-Nter (n = 18)** | 625 ± 4 | 0.62 ± 0.02 | 1.1 ± 0.03 | 3.3 ± 0.06 | 1.8 ± 0.04 | 0.082± 0.004 | 0.37 ± 0.01 | 84 ± 0.3 | 47 ± 0.4 |

**Table S2. Echocardiographic parameters of WT and *Scn5a*$^{+/-}$ mice injected or not with AAV-Nter.** Data are presented as mean ± SEM. Ctrl indicates control, WT wild type, HR Heart Rate, dPWT-LV diastolic Posterior Wall Thickness-Left Ventricle, sPWT-LV systolic Posterior Wall Thickness-Left Ventricle, dLVD diastolic Left Ventricle Diameter, sLVD systolic Left Ventricle Diameter, SV Stroke Volume, RWT relative wall thickness, EF Ejection Fraction and SF Shortening Fraction.

| | RR (ms) | P-wave duration (ms) | PR interval (ms) | QRS interval (ms) | QT interval (ms) |
|---|---|---|---|---|---|
| **Ctrl WT mice (n = 18)** | 184 ± 8 | 16.9 ± 0.4 | 40.4 ± 0.7 | 13.3 ± 0.3 | 54.4 ± 2.4 |
| **Ctrl *Scn5a*$^{+/-}$ mice (n = 19)** | 188 ± 9 | 22.9 ± 0.8**** | 46.8 ± 0.6 **** | 14.9 ± 0.4 ** | 54.9 ± 1.8 |
| ***Scn5a*$^{+/-}$ mice + AAV-Nter (n = 16)** | 191 ± 8 | 19.5 ± 0.8 ‡‡ | 43.6 ± 0.8 ‡‡ | 13.2 ± 0.2 ‡‡ | 53.9 ± 2.1 |

**Table S3. ECG parameters of WT and *Scn5a*$^{+/-}$ mice injected or not with AAV-Nter.** Data are presented as mean ± SEM. Ctrl indicates control and WT wild type. ***P*<0.01 and *****P*<0.0001 when compared to Ctrl WT mice (*One-way Anova test*). ‡‡ *P*<0.01 when compared to Ctrl *Scn5a*$^{+/-}$ mice (*One-way Anova test*).

| Type of cells | Peak current density (pA/pF) | Activation | | Inactivation | |
|---|---|---|---|---|---|
| | | $V_{1/2}$ (mV) | *k* (mV) | $V_{1/2}$ (mV) | *k* (mV) |
| **Ctrl iPSC-CM** | -97 ± 9 (n=12) | -38.5 ± 1.5 (n=12) | 4.7 ± 0.2 | -89.7 ± 3.1 (n=4) | 6.1 ± 0.5 |
| **Ctrl iPSC-CM +Ad-GFP** | -105 ± 7 (n=8) | -40.4 ± 1.7 (n=8) | 4.9 ± 0.3 | -88.4 ± 2.5 (n=7) | 5.5 ± 0.3 |

**Table S4. Kinetics properties of $I_{Na}$ recorded in iPSC-CMs.** Data are presented as mean ± SEM. Peak current density is given at -20 mV. Ctrl indicates control, $V_{1/2}$ half activation or inactivation value and *k* inverse slope factor.

| Assays | Antibody | Company Cat# and RRID | Dilutions |
|---|---|---|---|
| **Immunostaining** | | | |
| **Cardiomyocyte markers** | Rabbit anti-troponin T | Abcam Cat# ab45932 | 1/500 in 1/10 blocking solution |
| | Mouse anti-a-actinin | Sigma Aldrich Cat# A7811 | 1/500 in 1/10 blocking solution |
| **Viral transduction** | Rabbit anti-GFP | Torrey Pines Biolabs Cat# TP401 | 1/1000 in 1/10 blocking solution |
| **Nuclear staining** | DAPI | Sigma Aldrich Cat# D9542 | 1/1000 in 1/10 blocking solution |
| **Secondary antibodies** | Alexafluor 594 chicken anti-rabbit IgG | Thermofisher Scientific Cat# A21442 | 1/500 in 1/10 blocking solution |
| | Dylight 488 goat anti-mouse IgG | Bethyl Cat# A90-116D2 | 1/500 in 1/10 blocking solution |
| **Western blot** | | | |
| **Primary antibodies** | Rabbit anti-Nav1.5 | Alomone Labs Cat# ASC -005 | 1/200 in PBS-Tween 0.1% -Milk 5% |
| | Mouse anti-GAPDH | Proteintech Cat# 60004-1 -Ig | 1/20000 in PBS-Tween 0.1% -Milk 5% |
| | Mouse anti-flag | Sigma Aldrich Cat# F1804 | 1/500 in PBS-Tween 0.1% -Milk 5% |
| | Rabbit anti-N-cadherin | Proteintech Cat# 22018-1 -AP | 1/5000 in PBS-Tween 0.1% -Milk 5% |
| **Secondary antibodies** | Goat anti-mouse Dylight™ 800 | LicorBiosciences Cat# SA5 -35521 | 1/10000 in PBS-Tween 0.1% -Milk 5% |
| | Goat anti-rabbit Dylight™ 800 | LicorBiosciences Cat# SA5 -35571 | 1/10000 in PBS-Tween 0.1% -Milk 5% |
| | HRP-conjugated anti -mouse IgG | R&D System, #HAF007 | 1/10000 in PBS-Tween 0.1% -Milk 5% |
| | HRP-conjugated anti -rabbit IgG | R&D System, #HAF008 | 1/10000 in PBS-Tween 0.1% -Milk 5% |

**Table S5. Antibodies used in the study.**

| Assays | Target | Forward and Reverse primer sequences | qPCR conditions |
|---|---|---|---|
| Validation of viral transduction (qPCR) | Nter-flag | 5'-GAGGACCTGGACCCCTTCTA-3'<br>5'-TCCTCGAGTAAGAGCGAGTGA-5' | Tm 60°C; 40 cycles<br>167 pb |
| | eGFP | 5'-GCAGAAGAACGGCATCAAGGT-3'<br>5'-ACGAACTCCAGCAGGACCATG-5' | Tm 60°C; 40 cycles<br>204 pb |
| | Nter | 5'-CTGGACCCCTTCTATAGCACCC-3'<br>5'-GGACTGAGGACATACAAGGCGT-3' | Tm 60°C; 40 cycles<br>101 pb |
| House-keeping gene (qPCR) | hRPLP0 | 5'-CAACCCAGCTCTGGAGAAAC-3'<br>5'-AGCAGCTGGCACCTTATTG-3' | Tm 60°C; 40 cycles<br>118 pb |
| | mRplp0 | 5'-CAACCCAGCTCTGGAGAAAC-3'<br>5'-AGCAGCTGGCACCTTATTG-3' | Tm 60°C; 40 cycles<br>118 pb |
| Quantifying $Na_v1.5$ (qPCR) | $Na_v1.5$ | 5'-CCATCGCAGTGGCTGAGTC-3'<br>5'- CCACCAGACACAACTTGGGA-3' | Tm 60°C; 40 cycles<br>115 pb |

**Table S6. Sequences of primers used for quantitative-PCR analysis.**

### SUPPLEMENTAL FIGURES

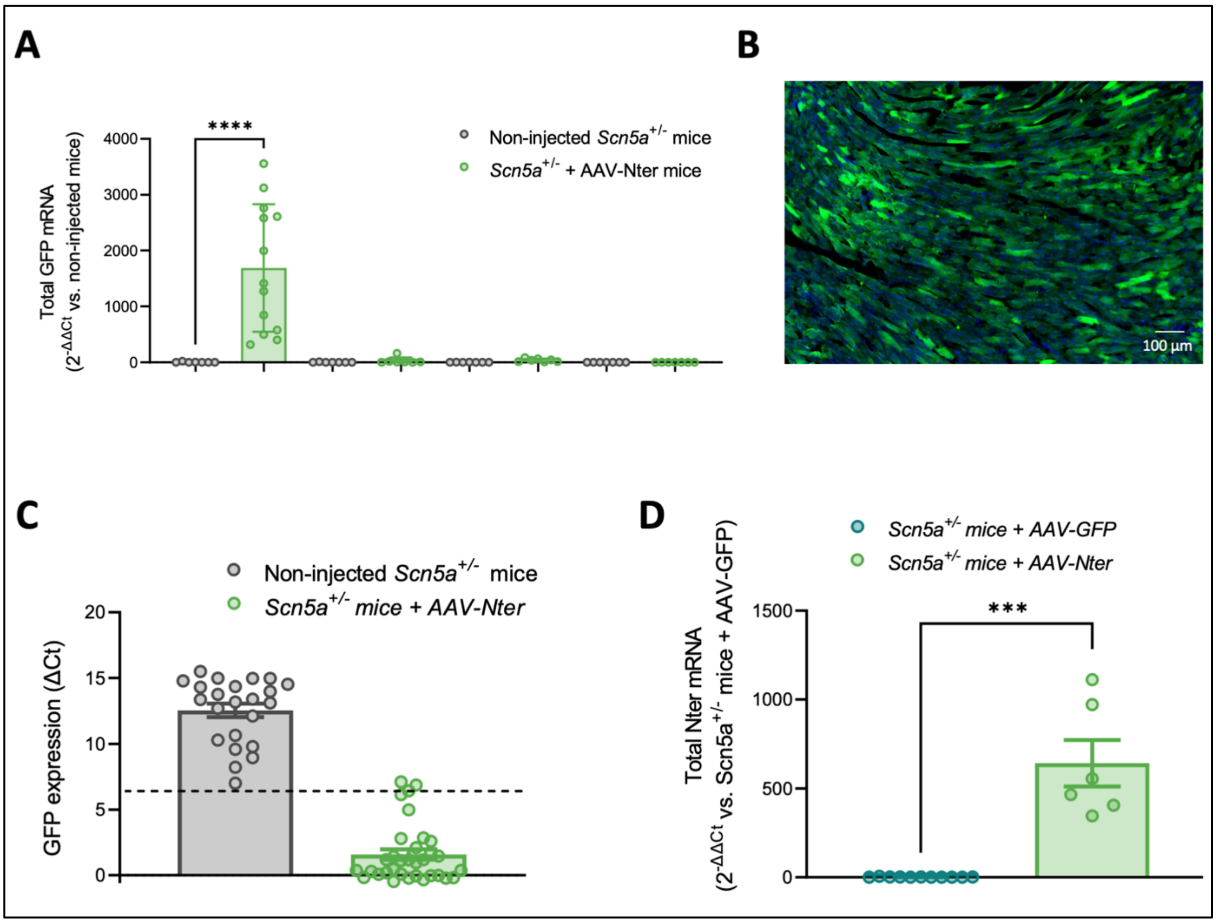


**Figure S1. AAV-mediated expression of Nter in mice is cardiac-specific. A.** GFP mRNA expression levels measured in heart, skeletal muscle, liver and brain tissues from non-injected *Scn5a*$^{+/-}$ mice or *Scn5a*$^{+/-}$ mice injected with AAV-Nter. GFP expression was normalized to *Rplp0* and the ratio AAV-Nter injected mice *vs.* non-injected mice ($2^{-\Delta\Delta Ct}$) was then calculated. Ct indicates Cycle threshold. ****$P<0.0001$ (*Anova test*). **B.** Representative image of 10-µm cryosections of mouse hearts injected with Nter showing GFP in green and nuclei in blue. **C.** AAV-Nter injection efficiency was validated by quantification of GFP mRNA expression normalized to *Rplp0*. GFP mRNA was quantified by $\Delta Ct = Ct^{GFP} - Ct^{Rplp0}$ in both *Scn5a*$^{+/-}$ mice and *Scn5a*$^{+/-}$ mice + AAV-Nter groups. The threshold was set at the lowest ΔCt value of the non-injected group and all injected mice with ΔCt values above this threshold were considered not-properly injected and excluded from the study. **D.** Total Nter mRNA expression level (exogenous and endogenous) measured in ventricular cardiac tissue of *Scn5a*$^{+/-}$ mice + AAV-GFP and *Scn5a*$^{+/-}$ mice + AAV-Nter. Cts were normalized to *Rplp0* and the ratio *Scn5a*$^{+/-}$ mice +

AAV-Nter *vs. Scn5a*$^{+/-}$ mice + AAV-GFP ($2^{-\Delta\Delta Ct}$) was then calculated; ****P*<0.001 (*Mann-Whitney test*).

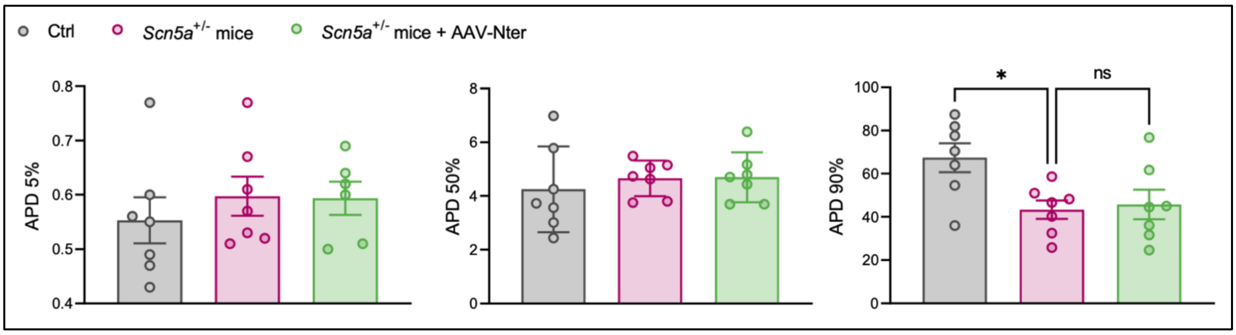


**Figure S2. Nter did not modify action potential durations in *Scn5a*$^{+/-}$ mice.** Action potential durations (APDs) at 5 ($APD_5$), 50 ($APD_{50}$) and 90 % ($APD_{90}$) of repolarization recorded in Ctrl, *Scn5a*$^{+/-}$ mice and *Scn5a*$^{+/-}$ mice + AAV-Nter; **P*<0.05 (*one-way Anova test*).

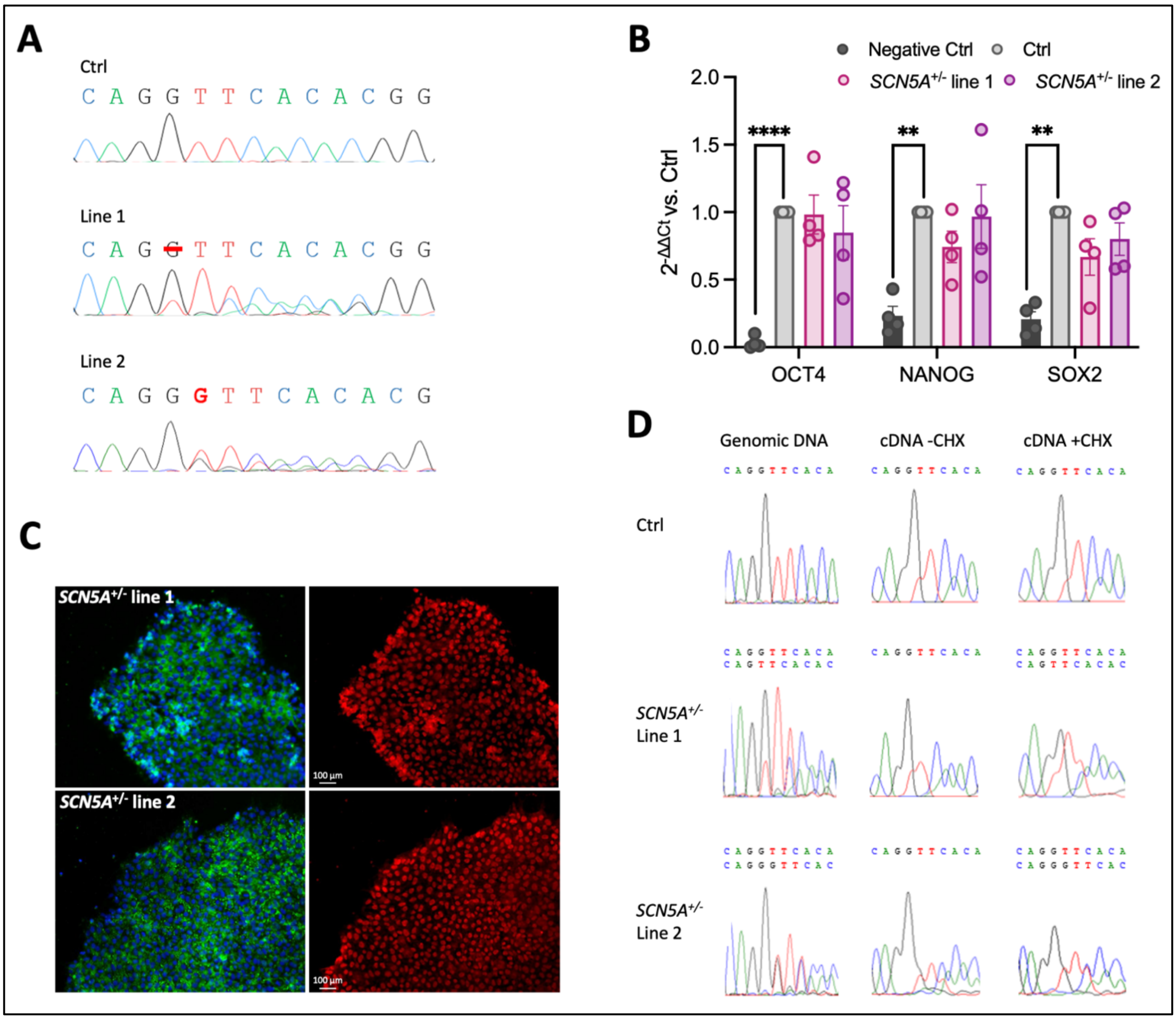


**Figure S3. Characterization of *SCN5A*$^{+/-}$ iPSCs. A.** Genomic sequence chromatograms validating *SCN5A* heterozygous mutation in CRISPR/Cas9 edited iPSC lines. **B.** Pluripotency marker (*OCT4*, *NANOG*, *SOX2*) mRNA expression level measured in cardiomyocytes of control (Ctrl) and *SCN5A*$^{+/-}$ iPSC. Cts were normalized to *RPLP0* and the ratio *vs.* Ctrl ($2^{-\Delta\Delta Ct}$) was then calculated; ***P*<0.01, *****P*<0.0001 (*two-way Anova test*). **C.** Immunostaining of pluripotency markers OCT-4 (red) and TRA1-81 (green) in both *SCN5A*$^{+/-}$ iPSC lines. **D.** Sanger sequencing chromatograms traces of cDNA from Ctrl and *SCN5A*$^{+/-}$ iPSC-CMs treated without (CHX-) or with (CHX+) cycloheximide, a blocker of translation. N=3 independent biological experiments per condition.

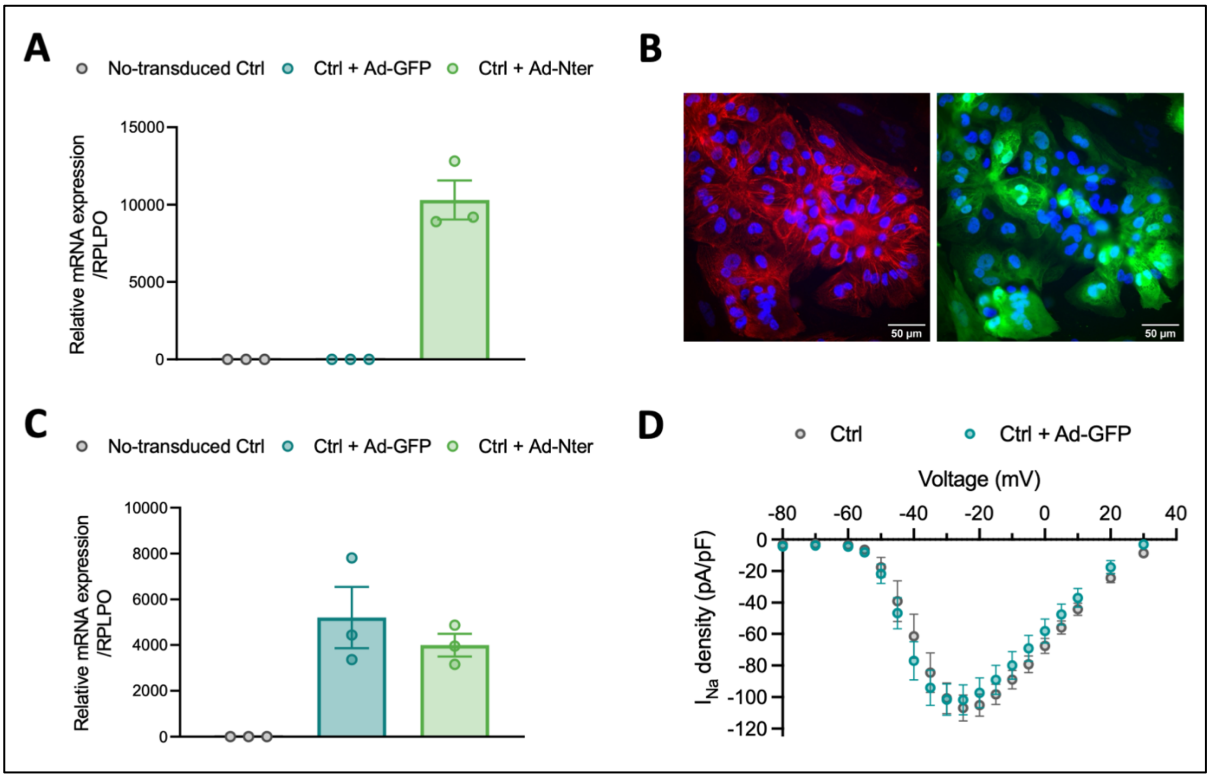


**Figure S4. Efficient transduction of iPSC-CMs by adenoviral vectors. A.** GFP mRNA expression level measured in Ctrl iPSC-CMs and *SCN5A*$^{+/-}$ iPSC-CMs transduced with Ad-GFP or Ad-Nter. Cts were normalized to *RPLP0* and the ratio *vs.* Ctrl ($2^{-\Delta\Delta Ct}$) was then calculated. **B.** Representative image of α-actinin (red), GFP (green) and nuclei (blue) immunostaining in iPSC-CMs. **C.** Nter-flag mRNA expression level measured in Ctrl iPSC-CMs and *SCN5A*$^{+/-}$ iPSC-CMs transduced with Ad-GFP or Ad-Nter. Cts were normalized to *RPLP0* and the ratio *vs.* Ctrl ($2^{-\Delta\Delta Ct}$) was then calculated. **D.** $I_{Na}$ density-membrane potential relationships recorded in non-transduced iPSC-CMs and iPSC-CMs + Ad-GFP. N=3 independent biological experiments per condition.

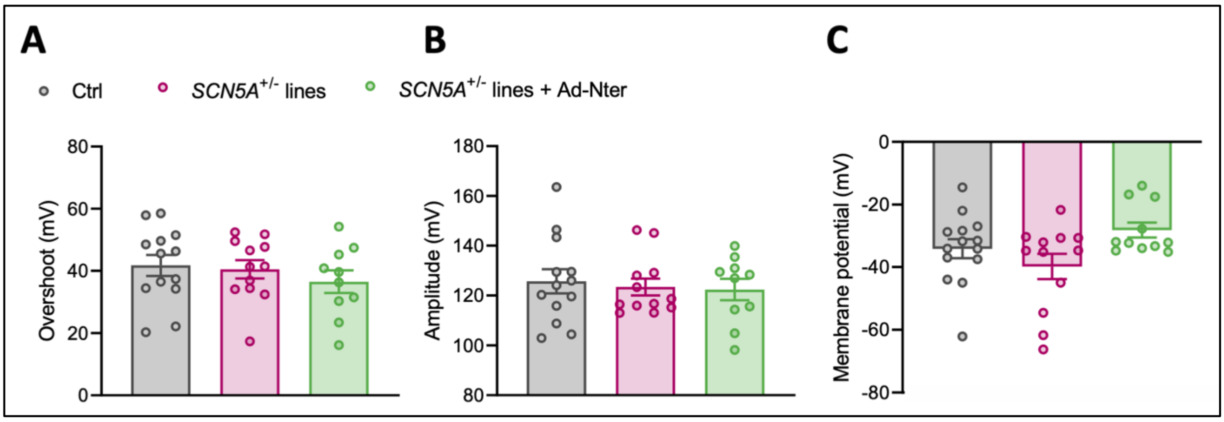


**Figure S5. AP parameters other than dV/dt$_{max}$ and duration were unaffected by *SCN5A* haploinsufficiency and Nter overexpression in iPSC-CMs. A.** Overshoot of action potentials (APs) recorded in control (Ctrl) iPSC-CMs and *SCN5A*$^{+/-}$ iPSC-CMs transduced with Ad-GFP or Ad-Nter. **B.** Amplitude of APs of Ctrl iPSC-CMs and *SCN5A*$^{+/-}$ iPSC-CMs transduced with Ad-GFP or Ad-Nter. **C.** Resting membrane potential of Ctrl iPSC-CMs and *SCN5A*$^{+/-}$ iPSC-CMs transduced with Ad-GFP or Ad-Nter. N=3 independent biological experiments per condition.
